# Arousal and locomotion differently modulate activity of somatostatin neurons across cortex

**DOI:** 10.1101/2022.01.18.476770

**Authors:** Christine F. Khoury, Noelle G. Fala, Caroline A. Runyan

## Abstract

Arousal powerfully influences cortical activity, in part by modulating local inhibitory circuits. Somatostatin-expressing inhibitory interneurons (SOM) are particularly well-situated to shape local population activity in response to shifts in arousal, yet the relationship between arousal state and SOM activity has not been characterized outside of sensory cortex. To determine whether SOM activity is similarly modulated by behavioral state across different levels of the cortical processing hierarchy, we compared the behavioral modulation of SOM neurons in auditory cortex (AC), a primary sensory region, and posterior parietal cortex (PPC), an association-level region of cortex. Behavioral state modulated activity differently in AC and PPC. In PPC, transitions to high arousal were accompanied by large increases in activity across the full PPC neural population, especially in SOM neurons. In AC, arousal transitions led to more subtle changes in overall activity, as individual SOM and Non-SOM neurons could be either positively or negatively modulated during transitions to high arousal states. The coding of sensory information in population activity was enhanced during periods of high arousal in AC, but not PPC. Our findings suggest unique relationships between activity in local circuits and arousal across cortex, which may be tailored to the roles of specific cortical regions in sensory processing or the control of behavior.

**Significance statement:** The effects of arousal on brain networks are profound, but vary across regions. Somatostatin neurons may carry out some of the effects of arousal on local network activity in sensory cortex, by modulating response gain and decorrelating population activity. However, SOM neurons have not been well studied outside of sensory cortex, and so it is unknown whether SOM neurons are similarly affected by shifts in brain state throughout the cortex. Here, we have revealed specialization in the relationship between arousal and activity in SOM neurons that could contribute to the diversity of arousal-related impacts on local computation across cortical regions.

## Introduction

Arousal profoundly impacts brain activity, at both local and global scales (Livingstone and Hubel, 1981; Marrocco et al., 1994; Cantero et al., 1999; Gould et al., 2011; Grent-’t-Jong et al., 2011; Stringer et al., 2019), and affects sensory perception by tailoring sensory processing to match behavioral demands (Pinto et al., 2013; McGinley et al., 2015; Kuchibhotla et al., 2016; Dadarlat and Stryker, 2017). In sensory cortex, the response gain and signal to noise ratio of individual neurons’ responses to sensory stimuli are enhanced with arousal (Niell and Stryker, 2010; Bennett et al., 2013; Polack et al., 2013; Fu et al., 2014; McGinley et al., 2015; Vinck et al., 2015; Mineault et al., 2016). At the population level, activity across neurons becomes decorrelated (Poulet and Petersen, 2008; Zhou et al., 2014; Vinck et al., 2015; Lin et al., 2019), which can enhance the encoding of sensory information by reducing redundancy and correlated noise (Averbeck et al., 2006).

In sensory cortex, these effects can be triggered by the release of neuromodulators linked to arousal, such as norepinephrine and acetylcholine, which act in part through inhibitory interneurons to influence local activity patterns (Fu et al., 2014; Chen et al., 2015; Kuchibhotla et al., 2016; Garcia-Junco-Clemente et al., 2019). The inhibitory cell class can be divided into three nonoverlapping subtypes, which express parvalbumin (PV), somatostatin (SOM), or vasoactive intestinal peptide (VIP), and exhibit distinct local connectivity patterns (Tremblay et al., 2016). SOM neurons are particularly sensitive to arousal state fluctuations, through direct activation of cholinergic and noradrenergic receptors (Kawaguchi and Shindou, 1998; Xiang et al., 1998; Beierlein et al., 2000; Fanselow et al., 2008; Chen et al., 2015; Kuchibhotla et al., 2016), and through inhibition by VIP neurons (Pfeffer et al., 2013; Pi et al., 2013; Fu et al., 2014; Karnani et al., 2016; Dipoppa et al., 2018). In turn, SOM neurons can powerfully influence local network activity, as they densely innervate the local excitatory population (Fino and Yuste, 2011; Pfeffer et al., 2013; Xu et al., 2013). When they are active, SOM neurons enhance stimulus selectivity and response reliability (Adesnik et al., 2012; Wilson et al., 2012; Rikhye et al., 2021), control local population activity dynamics (Chen et al., 2015; Veit et al., 2017), and flexibly modulate excitatory responses to stimuli, based on behavioral relevance (Kato et al., 2015; Wang and Yang, 2018).

The impacts of arousal on sensory perception, and on coding in sensory cortex, have been well studied (Fu et al., 2014; Zhou et al., 2014; McGinley et al., 2015; Vinck et al., 2015; Pakan et al., 2016). It is unclear though how specialized or generalized the relationship between arousal and local inhibitory circuit function is across the cortical hierarchy. For example, while arousal- and attention-mediated reductions in shared variability seem to improve sensory coding (Cohen and Maunsell, 2009; Goard and Dan, 2009), similar effects could be detrimental in the read-out of perceptual decisions to control behavior in higher cortex (Runyan et al., 2017; Valente et al., 2021). The basic structure of local circuits is highly conserved across cortical regions, yet differences in neuromodulatory receptor expression, the density of specific cell types, or in the specifics of local connectivity can alter the influence of neuromodulatory input on the pattern of neural population activity. Indeed, the density of SOM neurons is increased relative to PV neurons in association cortex (Kim et al., 2017; Dienel et al., 2020), suggesting that the population-level computations that SOM neurons participate in, and thus the relationship between arousal state and local network state, may differ across the cortical processing hierarchy.

Here, we hypothesized that arousal-related modulation of SOM neurons, and of local neural activity, would be specialized across cortical regions to match different arousal-related demands on local computation. We examined the effects of arousal state on activity in SOM and Non-SOM neurons in the primary auditory cortex (AC), and in the posterior parietal cortex (PPC). AC is a primary sensory region of the cortex, where the relationship between arousal and neural activity has been well studied (Schneider et al., 2014; Zhou et al., 2014; McGinley et al., 2015; Bigelow et al., 2019; Yavorska and Wehr, 2021). PPC is an association-level region that participates in flexible sensorimotor transformations (Fitzgerald et al., 2011; Harvey et al., 2012; Morcos and Harvey, 2016; Licata et al., 2017; Tseng et al., 2022). In PPC, task engagement is known to impact the structure of local population activity (Runyan et al., 2017; Pho et al., 2018; Valente et al., 2021), though relatively little is known about the specific contribution of generalized increases in arousal to PPC’s activity. Outside of a task context, firing rates of neurons in PPC are positively correlated with arousal level (Stitt et al., 2018), but the effects of arousal on specific inhibitory neuron types within PPC are not known. In the current study, we have revealed different relationships between arousal state, the structure of local population activity, and information coding in AC and PPC, suggesting that the effects of arousal on local processing are specialized across the cortical hierarchy.

## Materials and Methods

### 1.0 **Experimental Design and Statistical Analysis**

All pairwise comparisons were done with two-sided paired or unpaired permutation (i.e. randomization) tests with 10,000 iterations as indicated, where p<0.0001 indicates the highest significance achievable given the number of iterations performed. Given that the exact p value is unknown in these cases, p values of the highest significance are reported as such rather than as an exact value. All permutation tests were performed for differences in means. For statistical comparisons involving more than two groups, we used Kruskal-Wallis (non-parametric ANOVA) and used unpaired permutation tests post-hoc to determine which groups differed from each other. Data fell into natural groupings by 1) brain area (AC or PPC) and by 2) cell-type (SOM or Non-SOM), as indicated by expression of the red fluorophore, tdTomato. All bar plots show mean and bootstrapped 95% confidence intervals using 1000 iterations unless otherwise indicated. When multiple comparisons were made between groups, significance thresholds were Bonferroni-corrected. Sample sizes were chosen based on previous studies comparing population activity dynamics across brain areas or cell types (Runyan et al., 2017; Khan et al., 2018).

### 2.0 Animals

All procedures were approved by the University of Pittsburgh Institutional Animal Care and Use Committee. Homozygous SOM-Cre mice (Sst-IRES-Cre, JAX Stock #013044) were crossed with homozygous Ai14 mice (RCL-tdT-D, JAX Stock #007914) obtained from Jackson Laboratory, ME, USA, and all experiments were performed in the F1 generation, which expressed tdTomato in SOM+ neurons. Mice were group housed in cages with between 2 and 4 mice. Adult (8-24 weeks) male and female mice were used for experiments (4 male, 2 female). Mice were housed on a reversed 12 hr light/dark cycle, and all experiments were performed in the dark (active) phase.

#### 2.1 Surgical procedures

Mice were anesthetized with isoflurane (4% for induction, and 1-2% maintenance during surgery), and mounted on a stereotaxic frame (David Kopf Instruments, CA). Ophthalmic ointment was applied to cover the eyes (Henry Schein, NY). Dexamethasone was injected 12-24 hours prior to surgery, and carprofen and dexamethasone (Covetrus, ME) were injected subcutaneously immediately prior to surgery for pain management and to reduce the inflammatory response. Two 2 x 2 mm craniotomies were made over left AC and PPC (centered at 2 mm posterior and 1.75 mm lateral to bregma). For AC, the craniotomy was centered on the temporal ridge, and the posterior edge was aligned with the lamboid suture. 2mm biopsy punches were used to outline the circumference of the window before drilling.

1-4 evenly spaced ∼60 nl injections of the AAV1-synapsin-l-GCamp6f (Addgene, MA stock #100837) that had been diluted to a titer of ∼1×10^12 vg/mL using sterile PBS were made in each cranial window, centered in each craniotomy. A micromanipulator (QUAD, Sutter, CA) was used to target injections ∼250 μm under the dura at each site, where ∼60 nl virus was pressure-injected over 5-10 minutes. Pipettes were not removed until 5 minutes post-injection to prevent backflow. Dental cement (Parkell, NY) sealed a glass coverslip (3mm) over a drop of Kwik Sil (World Precision Instruments, FL) over the craniotomy. Using dental cement, a one-sided titanium headplate was attached to the right hemisphere of the skull. After mice had recovered from the anesthesia, they were returned to their home cages, and received oral carprofen tablets (Bio-Serv, NJ) for 3 days post-surgery.

### 3.0 Experimental Setup

#### 3.1 Two-photon microscope

Images were acquired using a resonant scanning two-photon microscope (Ultima Investigator, Bruker, WI) at a 30 Hz frame rate and 512 x 512 pixel resolution through a 16x water immersion lens (Nikon CF175, 16X/0.8 NA, NY). On separate days, either AC or PPC was imaged at a depth between 150 and 300 μm, corresponding to layers 2/3 of cortex. For AC imaging, the objective was rotated 35-45 degrees from vertical, and for PPC imaging, it was rotated to 5-15 degrees from vertical, matching the angle of the cranial window implant. Fields of view were 500 µm^2^ and contained 187±95 neurons, 20±10 (mean and standard deviation) of which were classified as SOM. Excitation light was provided by a femtosecond infrared laser (Insight X3, Spectra-Physics, CA) tuned to 920 nm. Green and red wavelengths were separated through a 565 nm lowpass filter before passing through bandpass filters (Chroma, ET525/70 and ET595/50, VT). PrairieView software (v5.5, Bruker, WI) was used to control the microscope.

#### 3.2 Behavioral monitoring

Running velocity was monitored on pitch and roll axes using two optical sensors (ADNS-98000, Tindie, CA) held adjacent to the spherical treadmill. A microcontroller (Teensy, 3.1, Adafruit, NY) communicated with the sensors, demixing their inputs to produce one output channel per rotational axis using custom code. Outputs controlling the galvanometers were synchronized with running velocity using a digital oscilloscope (Wavesurfer, Janelia, VA).

Pupil images were acquired at 1280 x 1024 pixels, at 10 Hz from an infrared (IR) camera focused on one eye (Flea3 FL3-U3-13Y3M-C ½” Monochrome USB 3.0 Camera, 1.0x SilverTL Telecentric Lens, FOV = 6.74mm X 5.39mm, Edmund Optics, NJ). The pupil was illuminated by the IR light emitted by the two-photon laser and required no additional IR illumination. Movies were acquired with the Matlab Image Acquisition Toolbox (Mathworks, MA). Pupil area was determined in each pupil movie frame post-hoc using custom Matlab code (Mathworks, MA). The pupil was constricted by controlling ambient illumination with an array of LCD screens (LG LP097QX1, South Korea), to maintain a moderate pupil area baseline from which increases and decreases in area could be measured.

#### 3.3 Experimental protocol

Imaging began 3-5 weeks post-surgery once robust expression of the GCaMP6f virus was observed. In each imaging session, GCaMP6f fluorescence changes were imaged in SOM (tdTomato+) and Non-SOM neurons, while mice ran freely on a spherical treadmill. In the spontaneous context, no sensory stimuli were delivered, while in the passive listening context, location-varying sound stimuli were presented (see Methods section 3.4). Spontaneous and passive listening contexts lasted ∼25-50 minutes each. Imaging alternated between AC and PPC across days. Multiple imaging sessions were performed in each cranial window, focusing at slightly different depths and lateral/posterior locations within the imaging windows across sessions. AC and PPC were each imaged in six mice (biological replicates). Each cranial window was imaged up to 11 times (technical replicates). Imaging from a given cranial window was suspended when we observed nuclear inclusion in 2 or more cells in the field of view, which indicates an over-expression of GCaMP6f.

#### 3.4 Sound stimuli

Four magnetic speakers were positioned in a semicircular array, centered on the mouse’s head (MF1-S, Tucker-Davis, FL). The speakers were positioned at -90, -30,+30 and +90 degrees from the midline in azimuth and driven by Matlab through a digital/analog converter (National Instruments). Speakers were calibrated to deliver similar sound levels (∼70 db) in a sound isolation chamber using a random incidence microphone (4939, Brüel & Kjær, Denmark). During passive listening, 1 or 2 second dynamic ripples (broadband stimuli created in Matlab by summing 32 tones spaced across 2-32 kHz, which fluctuated at 10-20 Hz (Elhilali et al., 2004) were presented from one of eight locations. Four of the sound locations corresponded to the locations of the four speakers (-90, -30, +30, +90 degrees), while the other four sound locations (-60, -15, +15, +60 degrees) were simulated using vector-based intensity panning, where the same sound stimulus was delivered to two neighboring speakers simultaneously, scaled by a gain factor (Runyan et al., 2017). Dynamic ripples were chosen to optimally drive populations of neurons in auditory cortex with diverse frequency tuning preferences. Each sound repeated three times at one location before switching to another. Each ripple played from each of the eight locations in randomized order, with a 240 ms gap between each sound. Output controlling the audio speakers recorded along with two-photon imaging galvo and running velocity using Wavesurfer (Janelia, VA), and these signals were aligned offline.

### 4.0 Data Processing

Imaging datasets from 24 AC fields of view and 20 PPC fields of view were included from 6 mice. We excluded any datasets with significant photobleaching or more than two filled cells. We also excluded any AC or PPC dataset from analysis if fewer than 1/3 of neurons that were significantly responsive (according to our definition in section 5.1) to at least one sound location, as we were interested in the effect of arousal on both spontaneous and sound-evoked responses. For AC datasets, we analyzed single-cell responses to pure tones on a subset of fields of view from each mouse, and then anatomically aligned all fields of view from datasets collected from each window, to ensure each field of view lay in a region representing tone frequencies in the sonic range of the tonotopic axis of primary auditory cortex. We eliminated any datasets where >50% of tone-responsive neurons had a preferred frequency was in the ultrasonic range (>20kHz), as well as any fields of view that were aligned anterior to a field of view where this was observed, to assure that we were seeing sound responses in the range of frequencies primarily represented by our dynamic ripples (described in section 3.4). We collected widefield fluorescence responses to pure tones in all AC cranial windows and observed pure tone responses in the sonic range for all AC windows; however, the extent of the viral expression within windows was too spatially limited to allow for mapping of specific regions.

#### 4.1 Image Processing

For each field of view, the raw calcium movies collected during the spontaneous activity and passive listening contexts were concatenated prior to motion correction, cell body identification, and fluorescence and neuropil extraction. These processing steps were performed using Suite2p 0.9.3 in Python (Pachitariu et al., 2017). Suite2p first registered images to eliminate brain motion, and clustered neighboring pixels with similar time courses into regions of interest (ROIs). ROIs were manually curated using the Suite2p GUI, to ensure that only cell bodies as opposed to dendritic processes were included in analysis, based on morphology. Cells expressing tdTomato (SOM cells), were identified using a threshold applied in the Suite2p GUI based on mean fluorescence in the red channel after bleed-through correction applied by Suite2p’s cell detection algorithm, along with manual correction. For each ROI, Suite2p returned a raw fluorescence timeseries, as well as an estimate of neuropil fluorescence that could contaminate the signal. For each cell, we scaled the neuropil fluorescence by a factor by 0.7 and subtracted this timeseries from the ROI’s raw fluorescence timeseries to obtain a neuropil-corrected fluorescence signal for each selected cell.

#### 4.2 ΔF/F and deconvolution

Once the neuropil corrected fluorescence was obtained for each neuron, we calculated ΔF/F for each cell in each frame by calculating (F-F_baseline_)/F_baseline_ for each frame, where F is the fluorescence of a given cell at that frame and F_baseline_ was the eighth percentile of that cell’s fluorescence spanning 450 frames before and after (∼15s each way, 30 s total). ΔF/F timeseries were then deconvolved to estimate the relative spike rate in each imaging frame using the OASIS toolbox (Friedrich et al., 2017). We used the AR1 FOOPSI algorithm and allowed the toolbox to optimize the convolution kernel, baseline fluorescence, and noise distribution. A threshold of 0.05 a.u. was applied to remove all events with low magnitude from deconvolved activity timeseries. All analyses were performed with both ΔF/F and deconvolved activity, and showed the same trends. Outside of Figure 1F and 1H only results using deconvolved activity are shown.

**Figure 1:**
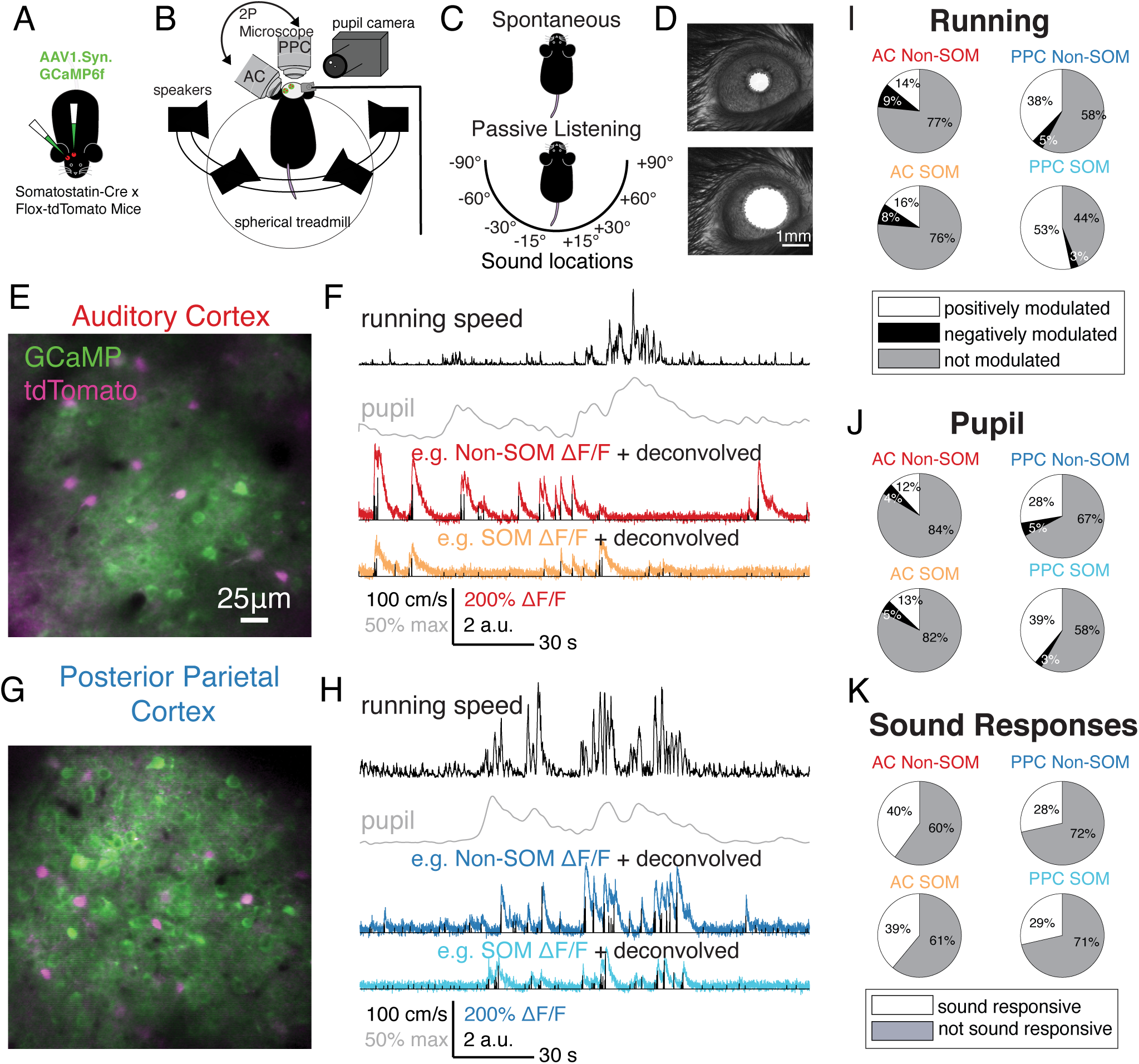
Imaging spike-related activity in SOM and Non-SOM neurons during behavioral state transitions. (A) Viral injections and cranial windows were made over auditory cortex (AC) and posterior parietal cortex (PPC) in each SOM-tdTomato mouse. (B) In imaging sessions, mice were headfixed over a spherical treadmill and allowed to run voluntarily. Four speakers arranged around the head presented sound stimuli. An infrared camera was used to image the pupil, and a rotating two-photon microscope was focused on either AC or PPC on a given imaging day. (C) Each imaging session included “Spontaneous” and “Passive Listening” contexts, without or with randomly presented sound stimuli from each of eight locations, respectively. (D) Pupil area was monitored via the pupil camera, scalebar in bottom image applies to top (constricted pupil) and bottom (dilated pupil). (E) Example field of view from auditory cortex, with intermingled tdTomato+/SOM+ (magenta) and tdTomato-/SOM-neurons, co-expressing GCaMP6f (green). (F) Example aligned behavioral and neural signals collected during the imaging session in E, including running speed (cm/s), normalized pupil area, dF/F from a Non-SOM (red) and SOM neuron (orange), each overlaid with the neuron’s deconvolved estimated spike rates. (G) As in E, for an example posterior parietal cortex field of view. (H) As in F, for the PPC field of view in G. (I) Proportions of Non-SOM and SOM neurons in AC (left) and PPC (right) with significant positive (white), negative (black), or no modulation (gray) by mouse’s running onset. AC Non-SOM N=2645; AC SOM N=359; PPC Non-SOM N=4719; PPC SOM N=525. (J) As in I, for pupil dilation. (K) Proportion of Non-SOM and SOM neurons in AC and PPC that were significantly sound responsive to at least one location (white) or not significantly sound responsive (gray).

### 5.0 Single cell modulation by sound stimuli, running behavior, and pupil size

#### 5.1 Sound responsiveness

Each neuron’s deconvolved activity was z-scored across its entire timeseries, and trial-averaged. For each sound location, we then calculated the sound-evoked response as the difference between the mean activity during the sound presentation and the mean activity in the 240 milliseconds prior to sound onset. We then compared the evoked sound responses to shuffled distributions, where each cell’s activity was shifted randomly by at least 5 seconds in time relative to sound location time series, and the sound-evoked response was recalculated. This was repeated 1000 times. A neuron was considered to be sound responsive if it had a sound-evoked response at least one sound location that was greater than the 97.5^th^ percentile of the shuffled distribution.

#### 5.2 Running bouts and modulation

Running bout onsets were defined as transitions in speed from below 10 cm/s to above 10 cm/s, and required that the mean running speed in the 1 s following the transition was three times greater than the 1 s prior to running bout onset, and that the mouse maintained a minimum speed of 15 cm/s for the following two seconds.

Running modulation was calculated as the difference in mean activity of a cell in the 1 s prior to running bout onset and the mean activity of a cell in the 3 s window following running bout onset. A shuffling procedure was applied to determine which cells were positively, negatively, and not modulated by running. Each cell’s activity was shifted randomly by at least 5 seconds in time relative to running speed timeseries, and for 1000 time-shifted iterations, running modulation was recalculated. Positively modulated neurons had positive running modulation values higher than the 97.5^th^ percentile of that cell’s shuffled distribution. Negatively modulated cells had negative running modulation values lower than the 2.5^th^ percentile of that cell’s shuffled distribution. All other cells were considered to be not modulated by running speed increases.

#### 5.3 Pupil dilation events and modulation

Pupil area was normalized to its maximum across the imaging session. To identify pupil dilation events, we first identified all local maxima of the pupil area. We then found the point prior to this where the derivative of pupil area was zero. We included events where the time from the inflection point to the local maximum was at least a 40% increase in pupil area and that the change from inflection point to local maximum was less than 1 second, and that the local maximum was at least 50% of the maximum total area by the pupil during that imaging session. We considered each inflection point to be the onset of dilation events.

To capture all pupil dilation related activity, which had a slower time course than running (Figure 4), we calculated pupil modulation for each neurons as the difference between the mean activity in the 1-s time window prior to dilation event onset and the mean activity in the 5-s time window following dilation event onset. We applied the same shuffling procedure as described for running modulation (5.2) to determine which neurons were positively, negatively, and not modulated by pupil dilation events.

### 6.0 Arousal States

#### 6.1 Defining low and high arousal states based on pupil area

K-means clustering was applied to the full pupil area timeseries, which included both spontaneous activity and passive listening contexts, to classify each pupil area measurement as low, transitional, or high arousal. Each pupil area timeseries was mean normalized using the following equation:

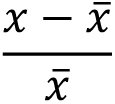

The Manhattan (aka City Blocks) distance metric was applied to define 2 centroid clusters that served as the high and low arousal groups. Transition periods included timepoints when the absolute difference in distance to the high and low arousal centroids was less than 0.05.

#### 6.2 Arousal Modulation Index

The arousal modulation index was calculated for each neuron using the following equation:

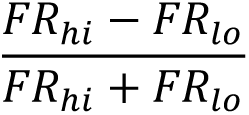

where FR_hi_ is the mean response of the neuron in the high arousal state and FR_lo_ is the mean response of the neuron in the low arousal state. We first maximum-normalized the deconvolved activity trace of each neuron across the entire time series. To calculate FR_hi_ (or FR_lo_), we summed the activity from the high (or low) arousal state and divided by the total time spent in the high (or low) arousal state in the spontaneous context. This index could vary continuously between -1 and +1, where negative values indicate higher activity in the low arousal state, and positive numbers indicate higher activity during the high arousal state.

### 7.0 Encoding Models

We used an encoding model to disentangle the contributions of pupil size and running speed to the activity of neurons in AC and PPC. In the Generalized Linear Model (GLM), the time-dependent effects of all measured external variables on each neuron’s activity were estimated (Pillow et al., 2008; Runyan et al., 2017). Three classes of predictors were used in different combinations to quantify their contributions to neuronal activity: running, pupil size, and sound stimulus predictors. We used a Bernoulli-based GLM to weight various combinations of predictors based on these variables to predict each neuron’s binarized activity (timeseries of relative spike rates were thresholded at 0.05). The encoding model is fully described in our previous work (Runyan et al., 2017).

#### 7.1 Pupil size and running predictors

Running velocity was measured at a higher time resolution than imaging and was binned to match the sampling rate of two-photon images (30 Hz). We included the velocity along the pitch and roll axes of the treadmill (relative to the mouse’s body axis). Running velocity measurements were separated into four channels, (1) forward, (2) reverse, (3) left, (4) right directions based on rotation along these axes. Running velocity changes could both precede or follow activity of individual neurons, so time series of running velocity were convolved with four evenly spaced Gaussian basis functions (240 ms half-width at half-height) extending 1s both forward and backward in time (8 basis functions total for each running direction: forward, reverse, left, and right). Changes in pupil area were modeled similarly. Because pupil area changes on a slower timescale, the pupil area trace was convolved with 16 evenly spaced Gaussian basis functions 4 s forward and backward in time to allow for either prediction or response to pupil area changes.

#### 7.2 Sound stimulus predictors

Sound stimuli were delivered from specific sound locations in the passive listening context. For sound stimulus onsets at each of the possible sound locations, 12 evenly spaced Gaussian basis functions (170 millisecond half width at half height) extended 2 s forward in time from each sound onset. First, second, and third repeats were represented separately due to potential adaptation-related effects. This resulted in 12 basis functions per repeat per sound location x 3 repeats x 8 locations for 288 sound predictors.

#### 7.3 GLM fitting and cross-validation procedures

All predictors were maximum normalized before the fitting procedure. Beta coefficients for the predictors were fitted to each neuron’s activity individually, using the glmnet package in R (Friedman et al., 2010) with elastic-net regularization, which smoothly interpolated between L_1_ and L_2_ type regularization according to the value of an interpolation parameter α, such that α=0 corresponded to L_2_ and α=1 corresponded to L_1_. We selected α=.25.

Trials were randomly split into training (70% of trials) and testing (remaining 30% of trials) sets, while balancing the distribution of sound locations. Fitting was performed on the training set, and within each training dataset, cross-validation folds (3x) were also pre-selected so that sound locations were evenly represented. Model performance (see below, 7.4) was assessed on the test set. Each model was thus fitted and tested on entirely separate data to prevent over-fitting from affecting results. This train/test procedure was repeated ten times, with random subsamples of the data included in train and test segments. Each model’s overall performance was assessed as its mean across all 10 iterations.

#### 7.4 GLM model performance

Each model’s performance for each cell was defined as the fraction of explained deviance of each model (compared to the null model). In the null model, only a constant (single parameter) was used to fit the neuron’s activity and no time-varying predictors were included. First, we calculated the deviance of the null and behavior model variants (see Methods section 7.5). For each model, the fraction of null model deviance explained by the model (d) was then calculated ((null deviance – model deviance)/null deviance). Deviance calculations were performed on a test dataset (30% of the data), which had not been included in the fitting procedure, and this train/test procedure was repeated ten times on randomly subsampled segments of the data.

#### 7.5 Running and pupil contribution

To identify the unique and separable contributions of running and pupil area on SOM and Non-SOM neurons’ activity, we fit three separate models: (1) full behavior model, (2) “no pupil” model, (3) “no running” model. In the full behavior model, all running, pupil, and sound predictors were included to fit each neuron’s activity. The “no pupil” model did not include the pupil predictors, and the “no running model” did not include the running predictors. Importantly, this analysis captures only the <u>unique</u> ways that pupil size and running can explain neural activity, where one cannot compensate for the other’s contribution.

We estimated the contribution of pupil or running to a neuron’s activity that could not be compensated for by the other variables, by comparing the model performance (fraction deviance explained, see Methods section 7.4) in the full vs. no pupil or no running models. The “running contribution” was calculated as the difference in fraction deviance explained of the full model and fraction deviance explained of the no-running model d_fb_-d_nr_. The “pupil contribution” was calculated as the difference in fraction deviance explained of the full model and fraction deviance explained of the no-pupil model d_fb_-d_np_.

### 8.0 Decoding

We used a population decoder to compare the sound location information contained in AC and PPC population activity, and its modulation with arousal state. The details of the decoder that we built to estimate the information about sound stimulus location have been previously described (Runyan et al., 2017). Briefly, for each trial we decoded sound stimulus location from single-trial population activity by computing the probability of external variables (sound location Left/Right category) given population activity. We used Bayes’ theorem, relying on population response probabilities estimated through the full behavior GLM and its predictors in that trial, to compute the posterior probability of each possible sound location stimulus. The decoder was “cumulative” in time, as for each time point t, it was based on all imaging frames from the initiation of the trial through time t. The decoded stimulus location at each time t was defined as the stimulus location with the maximum posterior probability, based on individual neurons or on a population of simultaneously imaged neurons. The population could include SOM, Non-SOM, or the “best” neurons. Non-SOM neurons were randomly subsampled 10 times, matching the sample size of SOM neurons in each iteration. The “best” neurons were selected as the n individual neurons with the best decoding performance, where n is the number of SOM neurons simultaneously imaged. Decoder performance was calculated as the fraction of correctly classified trials at each time point.

To compare decoder performance in low and high arousal states, trials were classified as “low” or “high” arousal based on normalized pupil area. Only the first sound repetition of each trial was used. Trials were randomly subsampled in the test set to ensure an even distribution of low and high arousal trials, and sound locations. This random subsample was repeated ten times.

### 9.0 Sound location sensitivity

To assess the location sensitivity of sound-related activity in SOM and Non-SOM neurons in AC and PPC, trial-averaged responses were used to calculate the “location sensitivity index”, based on vector averaging in the preferred sound direction:

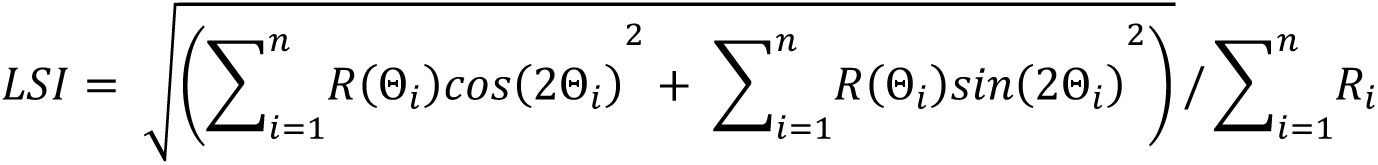

where R is the average response during the sound location presentation, and Θ is the sound location from -90 to +90 degrees, indexed by i = 1 to n (8 possible locations). LSI can vary continuously from 0 (unselective) to 1 (selectively responding to only one sound location). Datasets that did not have a minimum of 20 trials for each of the eight sound locations were excluded. Importantly, sounds were played from free-field speakers located at different angles relative to the interaural axis. A high LSI could correspond to true sound location selectivity based on interaural level differences, or to a neuron’s specific intensity tuning, due to different sound intensities impinging on the two ears.

### 11.0 Population-level Analyses

#### 11.1 Defining population activity axes related to sound location and arousal

To determine to what degree sound stimuli and arousal were driving population activity independently of each other in AC and PPC, we computed two axes for each imaging session: a sound location axis, and a pupil axis. The sound location axis was defined as the axis in population activity space that connects the mean response on -90° and +90° trials during the passive listening behavioral context, while the pupil axis was defined as the axis that connects the mean population activity during “low arousal” and “high arousal” periods during the spontaneous behavioral context, as defined by our pupil clustering algorithm. These definitions are analogous to how the “attention axis” is computed in primates (Cohen and Maunsell, 2010; Mayo et al., 2015; Cowley et al., 2020). We then measured the angle between the two vectors with a value from 0° to 90°, where 0° indicates linear dependence between the two subspaces and 90° indicates orthogonality.

#### 11.2 Computing Signal and Noise Correlations

We calculated noise correlations as fluctuations around mean sound responses, therefore we only included neurons that had significant sound responses (See: Figure 1K, Materials and Methods). Because it was rare for a mouse’s pupil to remain either in the high or low arousal cluster for the entire duration of a trial including all three sound repeats, we focused our analysis instead on the first repeat of a trial. We binned each neuron’s z-scored, deconvolved activity into 15-frame (∼500 ms) bins following sound onset. For each trial classified as high or low arousal, each cell’s mean sound response to all matching sound location presentations (including low, high, and unclassified trials) was subtracted and trials were concatenated. We computed partial Pearson correlations, discounting the effect of running speed (Matlab function ‘partialcorr’), on these traces. Because the ratio of low to high trials was variable across imaging sessions, we subsampled 10 times to balance for matching numbers of high and low trials at each sound location. We only considered imaging sessions in which there were at least 50 matched trials in low and high arousal clusters.

### 12.0 Histology

After all imaging sessions had been acquired, each mouse was transcardially perfused with saline and then 4% paraformaldehyde. The brain was extracted, cryoprotected, embedded, frozen, and sliced. Once slide mounted, we stained brains with DAPI to be able to identify structure. We used anatomical structure to verify the locations of our injections in AC and PPC.

## Results

To determine whether the effects of arousal on local activity are conserved across sensory and association cortex, we compared spontaneous and sensory-evoked activity in auditory cortex (AC) and posterior parietal cortex (PPC) in six mice of both sexes, during spontaneous shifts in the animals’ arousal state. We used two-photon calcium imaging in superficial cortex to measure the spike-related activity of neurons positive and negative for the red fluorophore tdTomato, which was expressed transgenically in somatostatin-positive (SOM) neurons (Madisen et al., 2009; Taniguchi et al., 2011). We virally expressed the genetically encoded calcium indicator GCaMP6f in all layer 2/3 neurons of AC and PPC in each mouse (Chen et al., 2013).

### Neural activity was modulated by sound stimuli and by behavioral correlates of arousal

During each imaging session, we focused the microscope on either AC or PPC, and imaged neural activity in two contexts. During the “spontaneous” context, the mouse ran freely on the spherical treadmill in the absence of sensory stimulation. During “passive listening”, sounds were presented from each of eight possible locations (Figure 1 B-C). In both contexts, mice were head-fixed and allowed to run voluntarily on a spherical treadmill. To track the mouse’s behavioral state, running velocity and pupil area were recorded throughout imaging sessions (Figure 1 A-H).

We first examined how changes in pupil area and running speed corresponded with changes in activity of individual neurons during the spontaneous context. Throughout imaging sessions, mice transitioned between behavioral states that reflect different arousal states: stillness and running, and pupil constriction and dilation (Figure 1 F, H). To quantify the effect of behavioral state transitions on individual neurons’ activity, we identified timepoints during the spontaneous context when either running speed or pupil area increased (Materials and Methods 5.2-5.3). The frequency of running bouts and dilation events was similar during PPC and AC imaging sessions (p=0.59 and p=0.65 respectively, permutation test, here and throughout unless otherwise noted, see Materials and Methods 1.0). Mice initiated running bouts at a rate of 0.915 bouts per minute [0.749, 1.11] (mean and [bootstrapped 95% confidence interval of the mean], here and throughout, unless otherwise indicated), and pupil dilations at 0.981 [0.857, 1.12] dilations per minute. We observed no difference between AC (N=24) and PPC (N=20) imaging sessions when considering pupil area and running speed during passive listening or spontaneous contexts (passive running speed: p=0.20; spontaneous running speed: p=0.52; passive pupil area: p=0.43; spontaneous pupil area: p=0.73). However, pupil area and running speed tended to be higher in general during the passive listening context (pupil: p=0.0020, running: p<0.0001, paired permutation test, N=44 datasets; Figure S1).

To determine how behavioral state affected neuronal activity, we next examined the activity of individual neurons during transitions from stationary to running, in the ‘spontaneous context’, when no sound stimuli were presented. We compared mean activity of each neuron (deconvolved estimated spike rates, here and throughout) in the 3-s time window following running bout onset to the 1-s time window prior to running bout onset (Figure 1 I, See Materials and Methods 5.2). Pupil size modulation was calculated similarly, based on transitions in pupil area from constricted to dilated (Figure 1 D, J; Materials and Methods, 5.3). We compared each neuron’s running and pupil modulation to shuffled distributions, where activity and behavioral data were time-shifted by random intervals, and classified each neuron as positively, negatively, or not modulated by behavioral state transition. Larger proportions of the SOM and Non-SOM populations were positively modulated by both pupil dilations and running bout onsets in PPC than in AC (Figure 1 I-J), suggesting that the effects of arousal on spontaneous activity are not uniform across areas. Among the groups considered, the PPC SOM population had the greatest proportion of neurons modulated by running speed and pupil dilation (Figure 1 I-J).

In the passive listening context, sound stimuli were presented from each of eight locations, centered on the mouse’s head (Figure 1 B-C, See Materials and Methods 3.4). We chose to manipulate sound location because of PPC’s role in spatial auditory processing during active behaviors (Nakamura, 1999). To determine whether each neuron was generally sound responsive, we computed the mean difference in activity during sound presentations and the pre-stimulus periods and compared to a shuffled distribution (Materials and Methods 5.1). We defined neurons as sound responsive if they responded to at least one sound location, more than would be expected from a random distribution obtained by shuffling. As expected, a greater proportion of AC SOM and Non-SOM neurons were sound responsive compared to PPC (39% of AC SOM and 40% of AC Non-SOM; 29% of PPC SOM and 28% of PPC Non-SOM neurons, Figure 1 K). The fraction of sound responsive neurons in AC is similar to the sparse sound encoding population described by others in layer 2/3 of AC (Hromádka et al., 2008). Furthermore, sound-evoked responses in AC Non-SOM neurons were lower when mice were running than when they were stationary (p<0.05, Figure S2), as has been reported on extensively by others (Schneider et al., 2014; Bigelow et al., 2019; Yavorska and Wehr, 2021). In PPC, sound-evoked responses were not affected by running behavior.

### Sound location coding in AC and PPC

To characterize the sound location sensitivity of responses in AC and PPC during the passive listening context, we computed each neuron’s sound location sensitivity index (LSI) (Figure 2 A-D, Materials and Methods 9.0). It is important to note that in the free-field sound stimulation configuration, we cannot distinguish true sound location selectivity from differences in sound intensity tuning. In both AC and PPC, Non-SOM neurons were more sensitive to sound location than SOM neurons (p<0.0001, Figure 2B-C). PPC neurons overall were less sensitive than AC neurons (Figure 2D), and PPC sound responses were less reliable than AC sound responses (p<0.001; Figure 2E). Interestingly, among neurons with sound location preferences, PPC sound responses were biased toward the lateral locations at +90 (contralateral) and -90 (ipsilateral) degrees, while the distribution of sound location preferences in AC neurons was more uniform (Figure 2F-G). Based on these differences in sound location sensitivity and response reliability, we expected population activity to encode sound location more accurately in AC than PPC.

**Figure 2:**
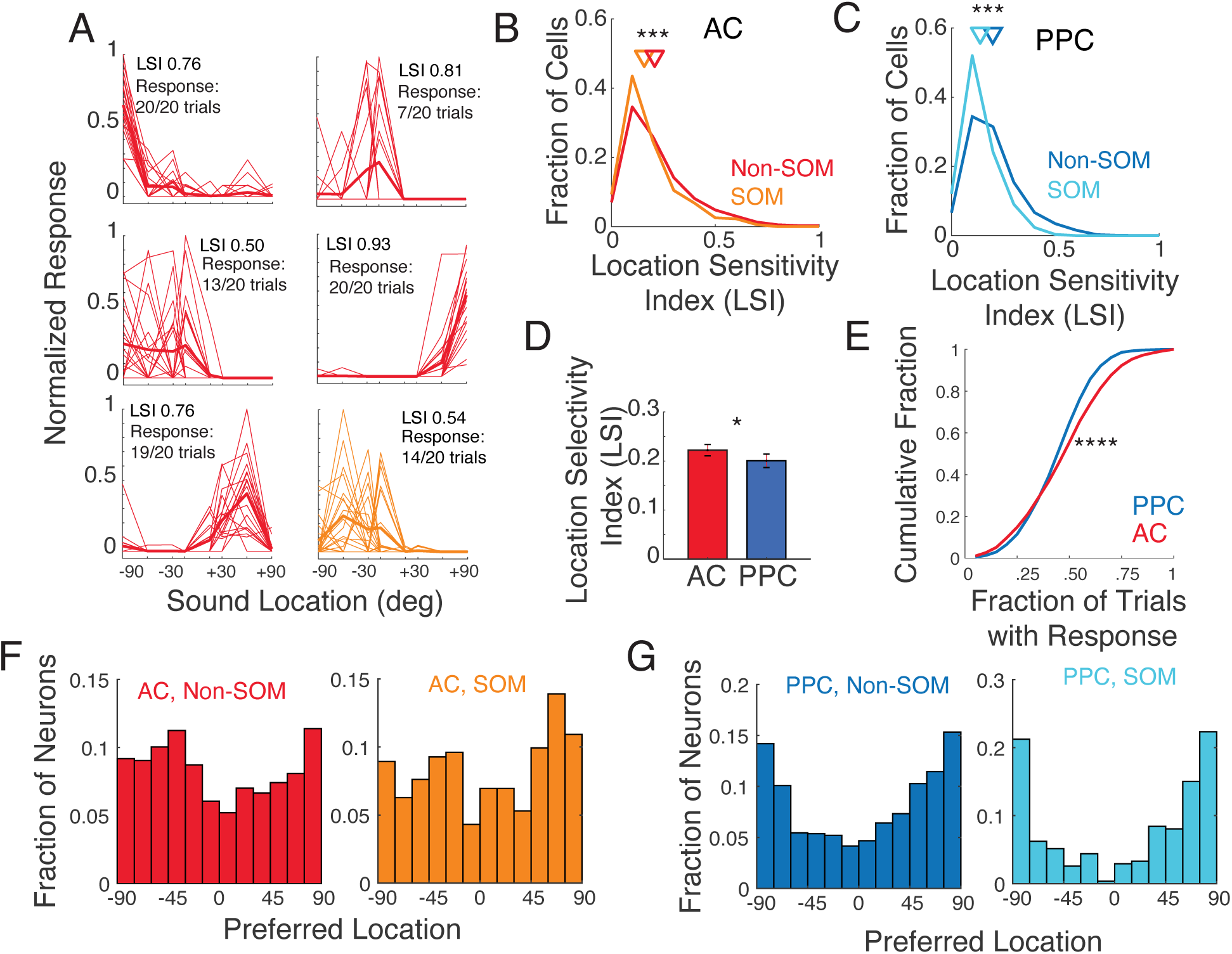
Sound Location Sensitivity in AC and PPC. (A) Six example neurons with diverse sound location preferences, ranging from ipsilateral (-90 - 0 degrees) to contralateral locations (0 - +90 degrees). Thin lines: mean response during sound presentation of individual trials. Thick lines: Trial average. The Location Sensitivity Index (LSI) and response reliability are reported for each example cell. Red: Example Non-SOM neurons. Orange: Example SOM neuron. (B) Distribution of Location Sensitivity Index (LSI) for Non-SOM (0.20 [0.20, 0.21], N=2455) and SOM (.19 [0.18, .19], N=354) neurons in AC. Triangles indicate population means. (C) As in B, for PPC Non-SOM (0.20 [0.19, 0.20], N=2553) and SOM (.13 [.12, .14], N=298). (D) Mean Location Sensitivity Index in AC and PPC. (E) Cumulative distributions of response reliability, the fraction of sound-responsive trials, in AC and PPC. (F) Histograms of the preferred locations of all AC Non-SOM and SOM neurons with LSI’s>0.05. (G) Histograms of the preferred locations of all PPC Non-SOM and SOM neurons with LSI’s>0.05. * p<0.05, ***p<0.001, ****p<0.0001, error bars indicate SEM.

To examine sound location coding at the level of neural populations, we constructed population decoders to predict the most likely sound stimulus location (left vs right) using the activity of different subsets of neurons. Each decoder was based on a Bayesian inversion of an encoding model that related each neuron’s activity to sound location and timing, running behavior, and pupil size (Figure 3A, Materials and Methods, 8.0; (Runyan et al., 2017)). The posterior probability of each stimulus location was computed cumulatively at each timepoint using population activity from all previous timepoints in the trial. Decoder performance was quantified as the fraction of trials where the stimulus with the maximal posterior probability matched the actual presented stimulus.

**Figure 3:**
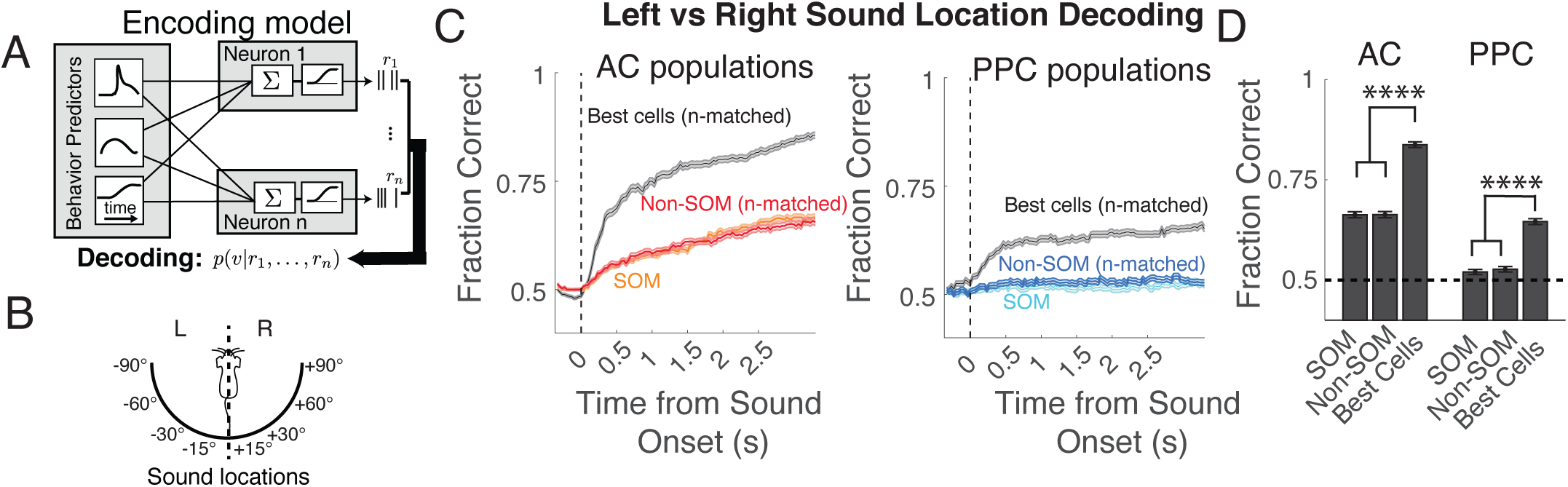
Decoding Sound Location from AC and PPC Population Activity. (A) The encoding model was trained on all trials that included all arousal levels, and inverted using Bayes Rule to compute the posterior probability of auditory stimuli given the activity of the neural population in AC and PPC. (B) Schematic of the discrimination being performed by the decoder, classifying sound stimuli as occuring from the left or right of the mouse. (C) Left: Mean cumulative population decoder performance (across datasets) when classifying left vs right locations, when based on “best cells”, defined as individual Non-SOM cells or SOM cells with the highest decoding performance. “Best cells” and Non-SOM cells in each dataset were subsampled to match the N of the SOM neuron population within each imaging field of view. Right: Population decoder performance using PPC neurons. Chance performance is 0.5. (D) Mean decoder performance using all timepoints in the trial (equivalent to the final timepoints in (C) using SOM neurons, subsampled Non-SOM neurons, and “best cells” in AC and PPC. Dotted line corresponds to chance performance (50%). N=24 AC datasets, N=20 PPC datasets. Error bars indicate SEM. ****p<0.0001.

First, to compare the overall ability of AC and PPC to represent sound location, we used activity of only the “best” cells (i.e., individual neurons whose activity best decoded sound location) regardless of cell type in the population decoder. The number of best cells used for each dataset was chosen to match the number of SOM neurons imaged during that session. The “best cell” decoder’s performance was above chance when using activity from both AC and PPC (Figure 3 C-D, Materials and Methods, 8.0). However, as expected based on the sound location sensitivity of individual neurons (Figure 2), AC decoding accuracy was higher than that of PPC (p<0.0001). Next, we compared the SOM and Non-SOM population decoders from each dataset. Within both areas, sound location decoding was similar when using the activity of either SOM or Non-SOM populations (AC: p=0.82; PPC: p=0.56, unpaired permutation tests, Figure 3 C), and both cell type populations from AC outperformed decoding based on PPC populations (p<0.0001, unpaired permutation test, Figure 3 C-D). All cell-type specific, subsampled population decoders performed worse than the decoders based on the n-matched population of “best” neurons (p<0.0001, paired permutation test; Figure 3 D, See Materials and Methods 8.0). To summarize, sound location was more accurately decoded from AC population activity than PPC, and SOM and random n-matched Non-SOM subpopulations in both areas were similarly informative about sound location. A sparse code for sound location was especially evident in AC, as small numbers of highly tuned “best” neurons more accurately encoded sound location than the random subsamples of the population. See Table 1 for full values and statistics.

**Table 1:**
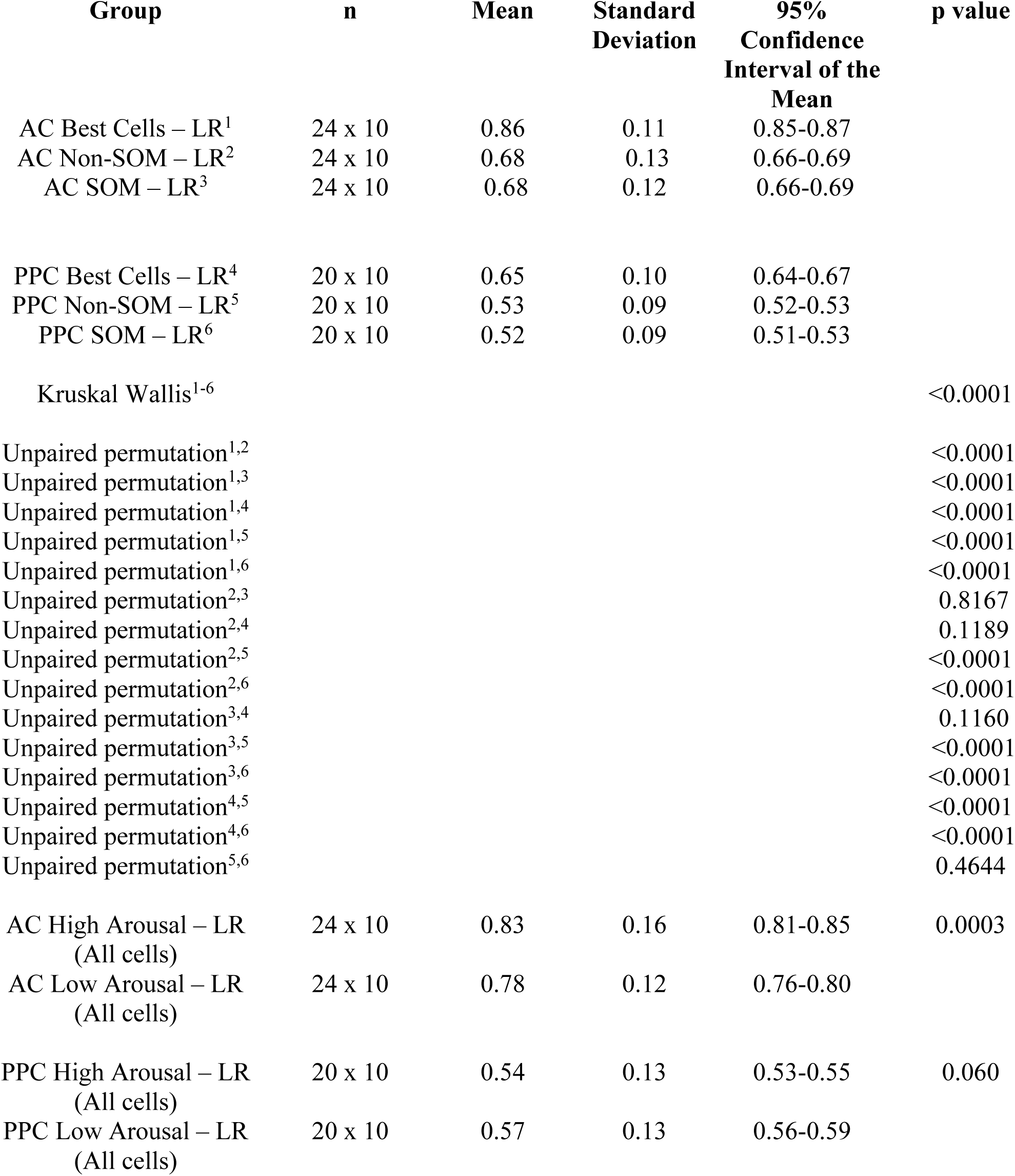
Sound Location Decoding/. Full values and statistics related to Figures 3 & 6

| **Group** | **n** | **Mean** | **Standard**<br>**Deviation** | **95%**<br>**Confidence**<br>**Interval of the**<br>**Mean** | **p value** |
| --- | --- | --- | --- | --- | --- |
| AC Best Cells – LR<sup>1</sup> | 24 x 10 | 0.86 | 0.11 | 0.85-0.87 |  |
| AC Non-SOM – LR<sup>2</sup> | 24 x 10 | 0.68 | 0.13 | 0.66-0.69 |  |
| AC SOM – LR<sup>3</sup> | 24 x 10 | 0.68 | 0.12 | 0.66-0.69 |  |
| PPC Best Cells – LR<sup>4</sup> | 20 x 10 | 0.65 | 0.10 | 0.64-0.67 |  |
| PPC Non-SOM – LR<sup>5</sup> | 20 x 10 | 0.53 | 0.09 | 0.52-0.53 |  |
| PPC SOM – LR<sup>6</sup> | 20 x 10 | 0.52 | 0.09 | 0.51-0.53 |  |
| Kruskal Wallis<sup>1-6</sup> |  |  |  |  | <0.0001 |
| Unpaired permutation<sup>1,2</sup> |  |  |  |  | <0.0001 |
| Unpaired permutation<sup>1,3</sup> |  |  |  |  | <0.0001 |
| Unpaired permutation<sup>1,4</sup> |  |  |  |  | <0.0001 |
| Unpaired permutation<sup>1,5</sup> |  |  |  |  | <0.0001 |
| Unpaired permutation<sup>1,6</sup> |  |  |  |  | <0.0001 |
| Unpaired permutation<sup>2,3</sup> |  |  |  |  | 0.8167 |
| Unpaired permutation<sup>2,4</sup> |  |  |  |  | 0.1189 |
| Unpaired permutation<sup>2,5</sup> |  |  |  |  | <0.0001 |
| Unpaired permutation<sup>2,6</sup> |  |  |  |  | <0.0001 |
| Unpaired permutation<sup>3,4</sup> |  |  |  |  | 0.1160 |
| Unpaired permutation<sup>3,5</sup> |  |  |  |  | <0.0001 |
| Unpaired permutation<sup>3,6</sup> |  |  |  |  | <0.0001 |
| Unpaired permutation<sup>4,5</sup> |  |  |  |  | <0.0001 |
| Unpaired permutation<sup>4,6</sup> |  |  |  |  | <0.0001 |
| Unpaired permutation<sup>5,6</sup> |  |  |  |  | 0.4644 |
| AC High Arousal – LR<br>(All cells) | 24 x 10 | 0.83 | 0.16 | 0.81-0.85 | 0.0003 |
| AC Low Arousal – LR<br>(All cells) | 24 x 10 | 0.78 | 0.12 | 0.76-0.80 |  |
| PPC High Arousal – LR<br>(All cells) | 20 x 10 | 0.54 | 0.13 | 0.53-0.55 | 0.060 |
| PPC Low Arousal – LR<br>(All cells) | 20 x 10 | 0.57 | 0.13 | 0.56-0.59 |  |

### SOM neurons’ activity was differently modulated during aroused states in AC and PPC

Next, we more thoroughly characterized the activity of SOM and Non-SOM neurons during arousal state transitions. We focused these analyses on the spontaneous context, to isolate the effects of state transitions from the effects of sound stimulation on activity. We defined behavioral state transitions as the onset times of “running bouts”, when the mouse’s running speed rapidly increased (Materials and Methods 5.2). We aligned and averaged the pupil area measurements to running bout onsets, observing that pupil area also increased during running bouts, though with a slower time course (Figure 4 A, gray trace, N=44 imaging sessions). We also aligned and averaged the activity of SOM and Non-SOM populations from AC and PPC to running bout onset (Figure 4 A, colored traces, AC, N=24; PPC, N=20 imaging sessions from 6 mice). In both AC and PPC, mean activity of SOM and Non-SOM neurons increased with the onset of running bouts. However, this increase in activity was weak in AC, due to the prevalence of both positively and negatively modulated SOM and Non-SOM neurons in the AC population (Figure 1I, Figure 4 B). In PPC, Non-SOM and SOM neurons more uniformly increased activity at running onset (Figure 1I, Figure 4 B).

**Figure 4:**
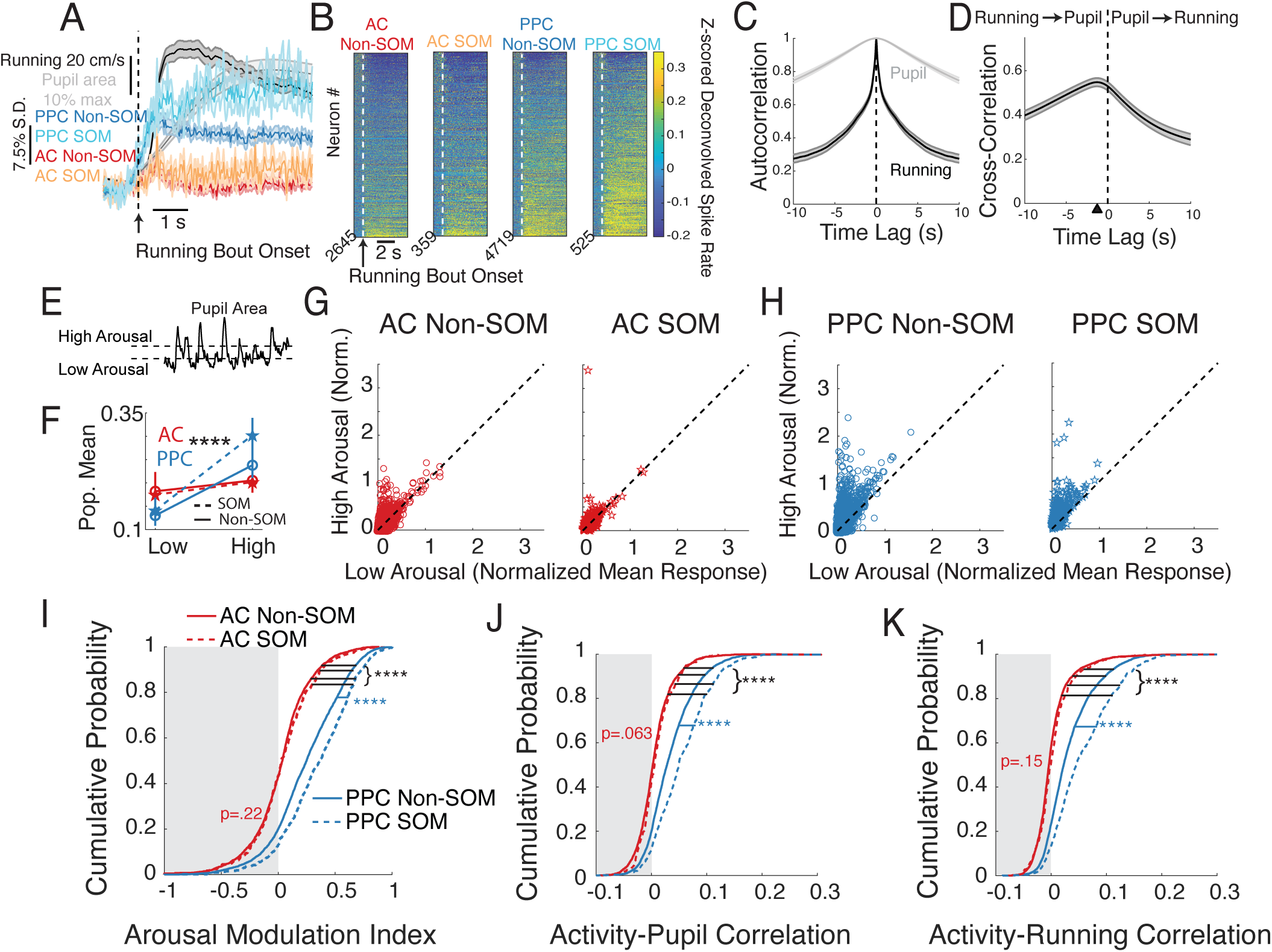
Single cell activity changes with arousal during the spontaneous context. (A) Average z-scored deconvolved activity, running speed, and pupil area aligned on running bout onset. Red: AC Non-SOM neurons’ activity (N=24 datasets), Orange: AC SOM neurons’ activity (N=24 datasets), Dark Blue: PPC Non-SOM neurons’ activity (N=20 datasets), Light Blue: PPC SOM neurons’ activity (N=20 datasets), Black: Running speed (N=44 datasets), Gray: Pupil area (N=44 datasets). (B) Average z-scored deconvolved activity of individual neurons, from L to R: all AC Non-SOM, AC SOM, PPC Non-SOM, and PPC SOM neurons aligned on running bout onset (AC Non-SOM N=2645; AC SOM N=359; PPC Non-SOM N=4719; PPC SOM N=525, in B-I). Neurons were sorted by running modulation. (C) Autocorrelation of pupil area and running speed, averaged across all datasets (N=44). (D) Cross-correlation between running speed and pupil area, averaged across all datasets (N=44). Black triangle indicates time lag with peak correlation, with running preceding pupil by 1.27 s. (E) Illustration of clustering that sorted pupil area during each imaging frame into low, transition, and high arousal states. (F) Mean responses of AC Non-SOM (red solid), AC SOM (red dotted), PPC Non-SOM (blue solid), and PPC SOM (blue dotted) neurons in low and high arousal states, as defined in E. (G) Left: Mean activity of AC Non-SOM neurons in high arousal state (classified with clustering as in E), plotted against mean activity in the low arousal state. Right: Mean activity of AC SOM neurons in high arousal vs low arousal states. (H) As in G, for PPC Non-SOM and SOM neurons. (I) Cumulative probability distribution of the arousal modulation index in AC Non-SOM (solid red), AC SOM (dotted red), PPC Non-SOM (solid blue), and PPC SOM (dotted blue) neurons. Arousal modulation index was calculated from the values in G-H, (High-Low)/(High+Low) for each neuron. (J) Cumulative probability distribution of the Pearson correlation between each neuron’s activity and pupil area, colors as in I. (K) Cumulative probability distribution of the Pearson correlation between each neuron’s activity and running speed. Significance as indicated, permutation test, **** p<.0001. Error bars: 95% bootstrapped confidence interval around the mean.

Because of the slow time course of changes in pupil area compared to the more rapid transitions in locomotion (Figure 4 C-D), we also characterized single neuron activity during periods of sustained pupil constriction and dilation. We classified pupil measurements as corresponding to low, transitional, or high arousal states (Figure 4 E, Materials and Methods 6.1), and focused analyses on the low and high arousal states. As expected from the relationship between running speed and pupil area, running speed was higher during the pupil-defined high arousal than low arousal states (p<0.0001). Mice ran on average 48.24±4.77 cm/s (mean±SEM) during the high arousal state, and 7.71±1.29 cm/s during the low arousal state (p<0.0001, N=44 datasets). SOM and Non-SOM activity was elevated overall in high arousal states (p<0.0001 for AC Non-SOM, PPC SOM and Non-SOM, p=0.0029 for AC SOM neurons, paired permutation test, Figure 4 E-H). To compare arousal modulation of activity in each population of neurons, we next computed an Arousal Modulation Index (AMI, Materials and Methods 6.2), which could vary from -1 to +1, with -1 indicating greater mean activity in the low arousal period, and +1 indicating greater activity in the high arousal period. The AMI was higher in PPC than AC neurons (p<0.0001, See Table 2 for full values and statistics). Interestingly, the AMI differed by cell type in PPC but not AC. In AC, the AMI was similar in SOM and Non-SOM neurons (p=0.22), while in PPC, the AMI was higher in SOM than Non-SOM neurons (p<0.0001, Figure 4 I).

**Table 2:** Arousal Modulation Index during spontaneous context. /Full values and statistics related to Figure 4I

| **Group** | **n** | **Mean** | **Standard**<br>**Deviation** | **95%**<br>**Confidence**<br>**Interval of the**<br>**Mean** | **p value** |
| --- | --- | --- | --- | --- | --- |
| AC SOM arousal modulation index<sup>1</sup> | 359 | 0.057 | 0.29 | 0.027-0.085 |  |
| AC Non-SOM, arousal modulation index<sup>2</sup> | 2645 | 0.035 | 0.28 | 0.025-0.046 |  |
| PPC Non-SOM arousal modulation index<sup>3</sup> | 4719 | 0.25 | 0.31 | 0.24-0.26 |  |
| PPC SOM arousal modulation index<sup>4</sup> | 525 | 0.35 | 0.31 | 0.32-0.37 |  |
| Kruskal Wallis<sup>1-4</sup> |  |  |  |  | 7.2E-12 |
| Unpaired permutation<sup>1,2</sup> |  |  |  |  | 0.22 |
| Unpaired permutation<sup>1,3</sup> |  |  |  |  | <0.0001 |
| Unpaired permutation<sup>2,3</sup> |  |  |  |  | <0.0001 |
| Unpaired permutation<sup>1,4</sup> |  |  |  |  | <0.0001 |
| Unpaired permutation<sup>2,4</sup> |  |  |  |  | <0.0001 |
| Unpaired permutation<sup>3,4</sup> |  |  |  |  | <0.0001 |

Finally, to consider the full time-varying relationship between ongoing neuronal activity and behavioral correlates of arousal, we correlated each neuron’s activity to pupil area and running speed, across the entire spontaneous behavioral context (Figure 4 J-K). Consistent with the above analyses based on activity aligned on state transitions, SOM and Non-SOM activity in PPC was more strongly correlated with both running and pupil size than in AC (p<0.0001). Within AC, SOM and Non-SOM activity was similarly correlated with the two behavioral measures (pupil area, p=0.063; running speed, p=0.146), while within PPC the activity of SOM neurons was again more strongly correlated with behavior than was the activity of Non-SOM neurons (p<0.001 for both running and pupil, Figure 4 J-K, See Table 3).

**Table 3:** Correlation coefficients with pupil area and running speed during spontaneous context. /Full values and statistics related to Figure 4J-K

| **Group** | **n** | **Mean** | **Standard**<br>**Deviation** | **95%**<br>**Confidence**<br>**Interval of the**<br>**Mean** | **p value** |
| --- | --- | --- | --- | --- | --- |
| AC Non-SOM, pupil correlation coefficient<sup>1</sup> | 2645 | 0.0057 | 0.033 | 0.0045-0.0070 |  |
| AC SOM pupil correlation coefficient<sup>2</sup> | 359 | 0.0091 | 0.031 | 0.0061-0.12 |  |
| PPC Non-SOM pupil correlation coefficient<sup>3</sup> | 4719 | 0.035 | 0.043 | 0.034-0.036 |  |
| PPC SOM pupil correlation coefficient<sup>4</sup> | 525 | 0.055 | 0.049 | 0.051-0.059 |  |
| Kruskal Wallis<sup>1-4</sup> |  |  |  |  | 6.3E-250 |
| Unpaired permutation<sup>1,2</sup> |  |  |  |  | 0.063 |
| Unpaired permutation<sup>1,3</sup> |  |  |  |  | <0.0001 |
| Unpaired permutation<sup>2,3</sup> |  |  |  |  | <0.0001 |
| Unpaired permutation<sup>1,4</sup> |  |  |  |  | <0.0001 |
| Unpaired permutation<sup>2,4</sup> |  |  |  |  | <0.0001 |
| Unpaired permutation<sup>3,4</sup> |  |  |  |  | <0.0001 |
| AC Non-SOM, running speed correlation coefficient<sup>1</sup> | 2645 | 0.0020 | 0.034 | 0.00060-0.0033 |  |
| AC SOM running speed correlation coefficient<sup>2</sup> | 359 | 0.0048 | 0.034 | 0.0013-0.0089 |  |
| PPC Non-SOM running speed correlation coefficient<sup>3</sup> | 4719 | 0.033 | 0.044 | 0.032-0.034 |  |
| PPC SOM running speed correlation coefficient<sup>4</sup> | 525 | 0.057 | 0.055 | 0.053-0.062 |  |
| Kruskal Wallis<sup>1-4</sup> |  |  |  |  | <0.0001 |
| Unpaired permutation<sup>1,2</sup> |  |  |  |  | 0.15 |
| Unpaired permutation<sup>1,3</sup> |  |  |  |  | <0.0001 |
| Unpaired permutation<sup>2,3</sup> |  |  |  |  | <0.0001 |
| Unpaired permutation<sup>1,4</sup> |  |  |  |  | <0.0001 |
| Unpaired permutation<sup>2,4</sup> |  |  |  |  | <0.0001 |
| Unpaired permutation<sup>3,4</sup> |  |  |  |  | <0.0001 |

Taken together, our results so far indicate that in the absence of sound stimulation, SOM and Non-SOM neurons have heterogeneous activity relationships with arousal state in AC, whether defined by running speed or pupil area. In PPC, neuronal activity was positively modulated with heightened arousal, with a stronger modulation of SOM neurons than Non-SOM neurons.

### An encoding model revealed different contributions of running speed and pupil size to single cell activity in AC and PPC

Running speed and pupil area are strongly correlated signals but vary on different timescales (Figure 4 C-D) and have separable effects on neuronal activity (Vinck et al., 2015). Our analyses of ongoing spontaneous activity in AC and PPC also hint at the possibility of separable effects of pupil area and running speed, as AC Non-SOM neurons’ activity was more highly correlated with pupil area than with running speed (p<0.0001, paired permutation test).

To disentangle the relationships between neuronal activity, running velocity, and pupil area, we used an encoding model approach (Pillow et al., 2008; Park et al., 2014; Runyan et al., 2017). We constructed a Generalized Linear Model (GLM) that used sound stimulus timing and location, pupil area, and running velocity to predict the responses of individual SOM and Non-SOM neurons in AC and PPC, in the passive listening context (Figure 5A). We note that in the above analyses (Figure 1, Figure 4), we considered only the speed at which the mouse was running in any direction, as increased running speed is correlated with heightened arousal (Fu et al., 2014; Zhou et al., 2014; Vinck et al., 2015; Mineault et al., 2016; Shimaoka et al., 2018). Because PPC neurons’ activity can be selective for running direction (Nitz, 2006; Whitlock et al., 2012; Runyan et al., 2017; Krumin et al., 2018; Minderer et al., 2019) in the GLM we used running velocity rather than speed to obtain more accurate predictions of each neuron’s activity (Materials and Methods 7.0, Figure 5).

**Figure 5:**
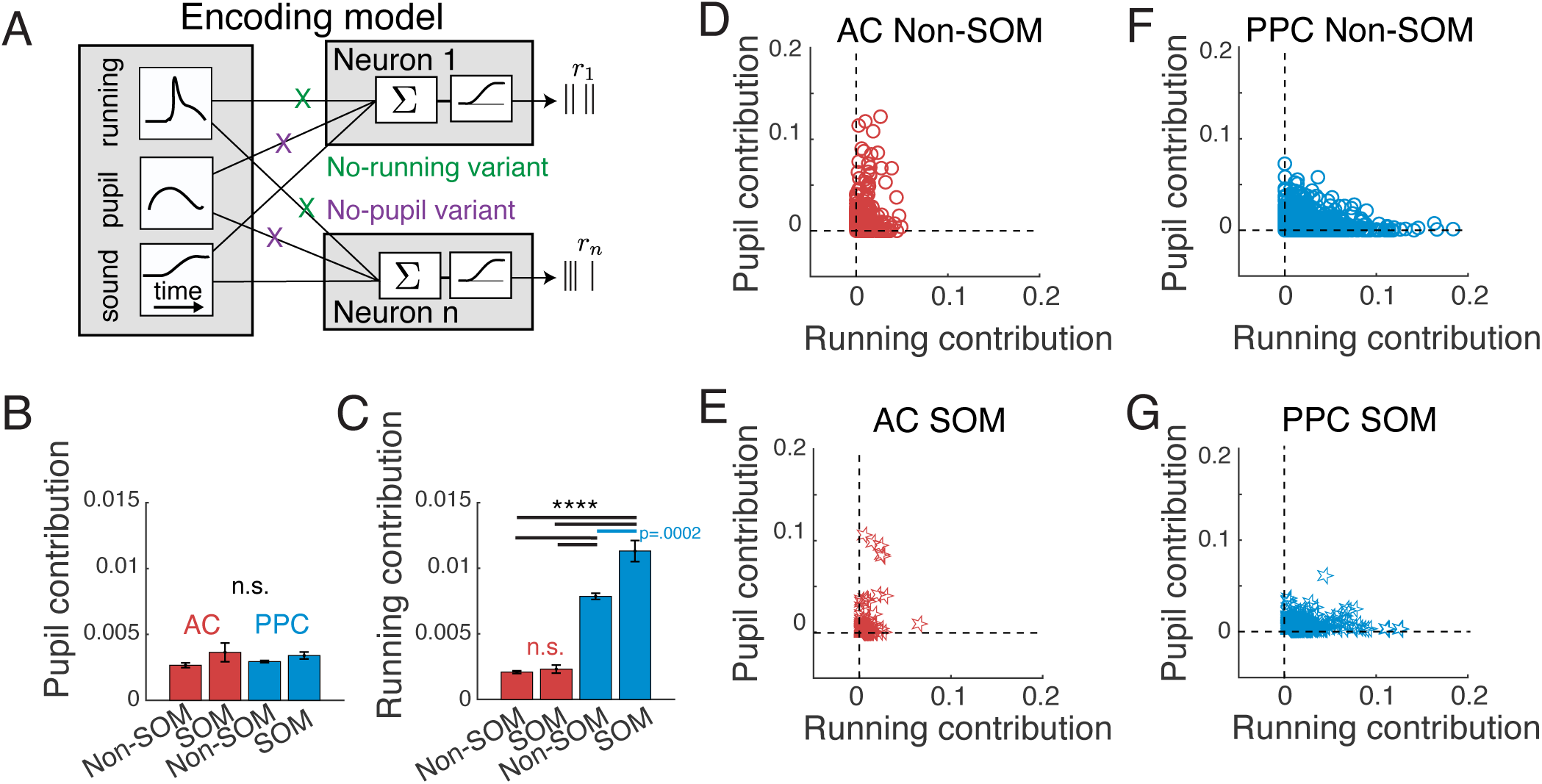
Using an Encoding Model to Disentange Running and Pupil Contributions to Single Cell Activity. (A) Schematic of Generalized Linear Model (GLM) used to determine the unique contributions of pupil area and running velocity to neurons’ activity. (B) Average contribution of pupil area to activity across all AC Non-SOM (N=2645) and SOM (N=359) neurons (red bars) and PPC Non-SOM (N=4525) and SOM (N=505) neurons (blue bars). Contribution was quantified as the improvement in model prediction performance of each neuron’s activity when adding pupil area as a set of predictors. Errorbars indicate SEM, n.s., not significant, exact values in text. (C) As in B, calculated when comparing model performance with and without running velocity predictors. AC Non-SOM and SOM neurons (red bars) and PPC Non-SOM and SOM neurons (blue bars), ****p<.0001. (D) Pupil contribution plotted against running contribution for each AC Non-SOM neuron, calculated as in B. (E) Pupil contribution plotted against running contribution for each AC SOM neuron. (F-G) Pupil contribution plotted against running contribution for each PPC Non-SOM or SOM neuron. Sample sizes in B apply to all panels.

To determine the relative contributions of pupil area and running velocity to neurons’ activity, we separately removed these predictors from the model and measured the decrement in the model’s prediction performance. For example, we calculated the pupil size contribution to a given neuron’s activity as the difference between the prediction performance of the model with and without the pupil size predictors. We considered this decrement in model performance as the contribution of pupil size to the neuron’s activity that is not redundant with running speed (Materials and Methods 7.5). If running velocity and pupil area did not make unique contributions to neuronal activity, running predictors would be able to account for the missing pupil area predictors and vice versa, and the performance of the model would not be degraded compared to the full model that includes both pupil area and running velocity predictors. The model comparison revealed single neuron activity that could be explained distinctly by running and by pupil in both AC and PPC (Figure 5 B-G, note neurons along the pupil and running contribution axes in D-G). While a Kruskal Wallis test indicated that the contribution of pupil size differed among the four cell-type/area combinations, post-hoc tests returned p>.05 for all pairwise comparisons (See Table 6 for full values and statistics). In contrast, running contributions were overall stronger in PPC neurons than in AC neurons (p<0.0001), and within PPC, running contributions were stronger to SOM than Non-SOM activity (p<0.001, Figure 5C, F, G). Within AC, running contributions did not depend on cell type (p=0.44, Figure 5C, D, E). See Table 6 for full values and statistics. Thus, running contributions, but not pupil contributions, reflect the observed differences in the arousal-dependence of activity in AC and PPC.

**Table 4:**
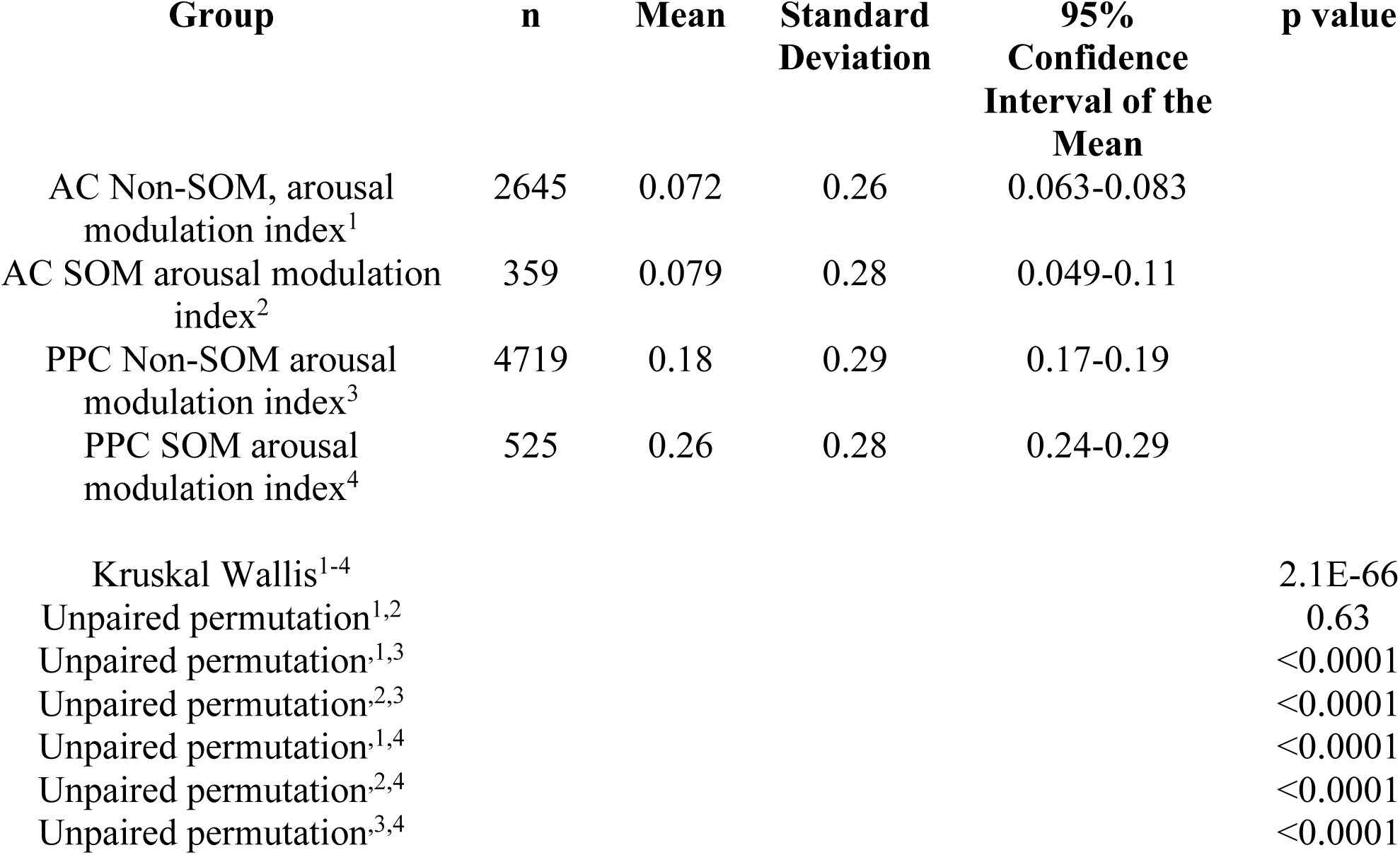
Arousal Modulation Index during passive listening context.

**Table 5:** Correlation coefficients with pupil area and running speed during passive listening context.

| **Group** | **n** | **Mean** | **Standard Deviation** | **95% Confidence Interval of the Mean** | **p value** |
| --- | --- | --- | --- | --- | --- |
| AC Non-SOM, pupil correlation coefficient<sup>1</sup> | 2645 | 0.0096 | 0.031 | 0.0081-0.010 |  |
| AC SOM pupil correlation coefficient<sup>2</sup> | 359 | 0.010 | 0.033 | 0.0074-0.014 |  |
| PPC Non-SOM pupil correlation coefficient<sup>3</sup> | 4719 | 0.030 | 0.042 | 0.029-0.031 |  |
| PPC SOM pupil correlation coefficient<sup>4</sup> | 525 | 0.043 | 0.044 | 0.038-0.046 |  |
| Kruskal Wallis<sup>1-4</sup> |  |  |  |  | 5.3E-115 |
| Unpaired permutation<sup>1,2</sup> |  |  |  |  | 0.63 |
| Unpaired permutation<sup>1,3</sup> |  |  |  |  | <0.0001 |
| Unpaired permutation<sup>2,3</sup> |  |  |  |  | <0.0001 |
| Unpaired permutation<sup>1,4</sup> |  |  |  |  | <0.0001 |
| Unpaired permutation<sup>2,4</sup> |  |  |  |  | <0.0001 |
| Unpaired permutation<sup>3,4</sup> |  |  |  |  | <0.0001 |
| AC Non-SOM, running speed correlation coefficient<sup>1</sup> | 2645 | 0.0067 | 0.029 | 0.0056-0.0078 |  |
| AC SOM running speed correlation coefficient<sup>2</sup> | 359 | 0.0076 | 0.30 | 0.0050-0.011 |  |
| PPC Non-SOM running speed correlation coefficient<sup>3</sup> | 4719 | 0.029 | 0.042 | 0.028-0.030 |  |
| PPC SOM running speed correlation coefficient<sup>4</sup> | 525 | 0.045 | 0.047 | 0.038-0.046 |  |
| Kruskal Wallis<sup>1-4</sup> |  |  |  |  | 1.2E-146 |
| Unpaired permutation<sup>1,2</sup> |  |  |  |  | 0.59 |
| Unpaired permutation<sup>1,3</sup> |  |  |  |  | <0.0001 |
| Unpaired permutation<sup>2,3</sup> |  |  |  |  | <0.0001 |
| Unpaired permutation<sup>1,4</sup> |  |  |  |  | <0.0001 |
| Unpaired permutation<sup>2,4</sup> |  |  |  |  | <0.0001 |
| Unpaired permutation<sup>3,4</sup> |  |  |  |  | <0.0001 |

**Table 6:** Pupil and Running Contributions/. Full values and statistics related to Figure 5

| **Group** | **n** | **Mean** | **Standard**<br>**Deviation** | **95%**<br>**Confidence**<br>**Interval of**<br>**the Mean** | **p value** |
| --- | --- | --- | --- | --- | --- |
| AC Non-SOM, pupil<br>diameter contribution<sup>1</sup> | 2476 | 0.0027 | 0.0086 | 0.0023-0.0031 |  |
| AC SOM, pupil diameter<br>contribution<sup>2</sup> | 330 | 0.0036 | 0.0123 | 0.0025-0.0049 |  |
| PPC Non-SOM, pupil<br>diameter contribution<sup>3</sup> | 4525 | 0.0029 | 0.0060 | 0.0028-0.0031 |  |
| PPC SOM, pupil diameter<br>contribution<sup>4</sup> | 505 | 0.0034 | 0.0059 | 0.0029-.0040 |  |
| Kruskal Wallis<sup>1-4</sup> |  |  |  |  | <0.0001 |
| Unpaired permutation<sup>1,2</sup> |  |  |  |  | 0.0780 |
| Unpaired permutation<sup>1,3</sup> |  |  |  |  | 0.1259 |
| Unpaired permutation<sup>2,3</sup> |  |  |  |  | 0.2360 |
| Unpaired permutation<sup>1,4</sup> |  |  |  |  | 0.0619 |
| Unpaired permutation<sup>2,4</sup> |  |  |  |  | 0.9520 |
| Unpaired permutation<sup>3,4</sup> |  |  |  |  | 0.1070 |
| AC Non-SOM, running<br>speed contribution<sup>1</sup> | 2476 | 0.0020 | 0.0050 | 0.0019-0.0023 |  |
| AC SOM running speed<br>contribution<sup>2</sup> | 330 | 0.0023 | 0.0005 | 0.0019-0.0320 |  |
| PPC Non-SOM running<br>speed contribution<sup>3</sup> | 4525 | 0.0079 | 0.0156 | 0.0075-0.0083 |  |
| PPC SOM running speed<br>contribution<sup>4</sup> | 505 | 0.0113 | 0.0181 | 0.0099-0.0131 |  |
| Kruskal Wallis<sup>1-4</sup> |  |  |  |  | <0.0001 |
| Unpaired permutation<sup>1,2</sup> |  |  |  |  | 0.4409 |
| Unpaired permutation<sup>1,3</sup> |  |  |  |  | 0.0001 |
| Unpaired permutation<sup>2,3</sup> |  |  |  |  | 0.0001 |
| Unpaired permutation<sup>1,4</sup> |  |  |  |  | 0.0001 |
| Unpaired permutation<sup>2,4</sup> |  |  |  |  | 0.0001 |
| Unpaired permutation<sup>3,4</sup> |  |  |  |  | 0.0002 |

### Sound location coding was enhanced with arousal in AC, but not PPC

Given the activity changes with arousal in AC and PPC, we hypothesized that arousal would affect sensory information coding in both regions. To test this possibility, we trained and tested the sound location decoder (Figure 3) in low and high arousal periods (Figure 6 A - E). Sound stimulus trials were classified as occurring during low or high arousal states based on pupil size (as in Figure 4 E), and low and high arousal trials were evenly balanced to train and test the decoders. Decoding performance for left vs. right sound locations using AC populations was modestly improved during heightened arousal (p=0.0003, paired permutation test), but decoding performance was similarly poor in low and high arousal trials using PPC population activity (Figure 6 C-E) (p=0.06, paired permutation test). Thus, while sensory coding in AC was slightly improved with heightened arousal, coding within PPC populations was unaffected by arousal state. See Table 1 for full values and statistics.

**Figure 6:**
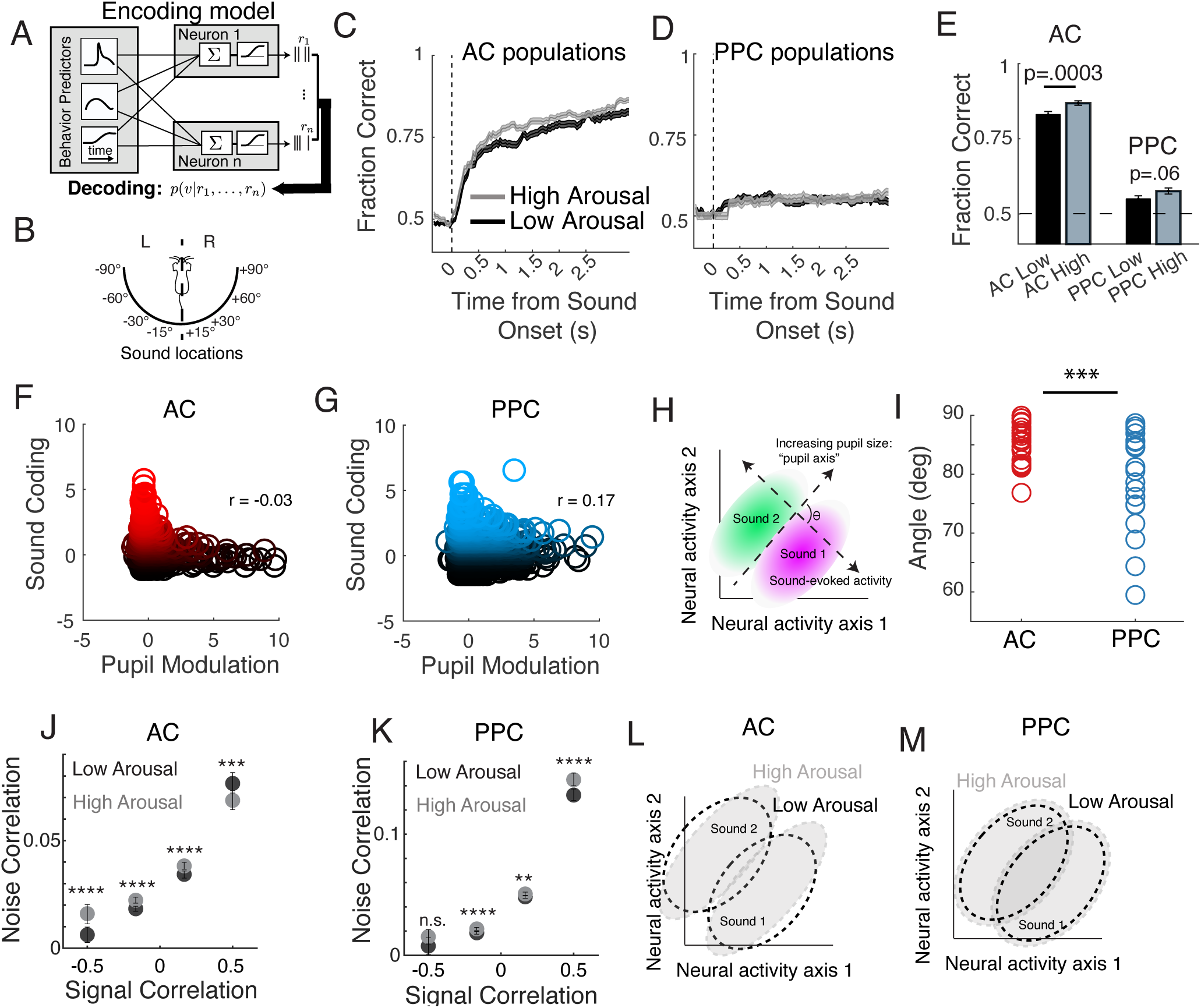
Arousal impacts the structure of population activity, improving information coding in AC. (A) The encoding model was trained on all trials that included all arousal levels, and inverted using Bayes Rule to compute the posterior probability of auditory stimuli given the activity of the neural population in AC and PPC. (B) Schematic of the discrimination being performed by the decoder, classifying sound stimuli as occuring from the left or right of the mouse. (C) Mean fraction correct of cumulative sound location (left sound locations vs right sound locations) decoding in AC (right) and PPC (left) populations in high (dark) and low (light) arousal conditions. Subsampling to match arousal and sound locations was repeated 10x. (D) Cumulative fraction correct of decoding for left vs right sound location at the end of the trial in AC and PPC. (F) For each AC neuron, the sound location decoding performance based on that neuron’s activity is plotted against that neuron’s pupil modulation, as quantified by the encoding model in Figure 5. Neurons are color-coded from black to red to indicate their z-scored sound location decoding performance (red neurons > 0 z-scored performance). (G) As in F, for PPC. (H) Schematic to explain the definitions of the sound location and pupil axes in high dimensional population activity space in I. (I) Angle in degrees between the pupil and sound location axes in each AC and PPC dataset. The angle between the pupil and sound axes is significantly smaller in PPC than AC. (J) Mean pairwise noise correlation among sound responsive AC neurons during high arousal (gray) and low arousal (black) trials, binned by their pairwise signal correlation. (K) As in (J), but for PPC. (L) Schematic demonstrating the impacts of arousal on sound coding in AC population activity. The shape of the population activity subspace responding to “sound 1” and “sound 2” changes between low arousal (dashed outlines) and high arousal (gray ovals) states, improving their discriminability. This shape change occurs because the shared variability is reduced among neurons coding for similar locations, and enhanced among neurons coding for different locations (see J). (M) As in L, for PPC. In PPC, the shape of the area of population activity space encoding sound 1 or sound 2 does not change with arousal, as shared variability is more generally increased across neuron pairs (see K). Across panels, ** p<0.01, *** p<0.001, **** p<0.0001, n.s. not significant. AC Non-SOM N=24 datasets; AC SOM N=24 datasets; PPC Non-SOM N=20 datasets; PPC SOM N=20 datasets throughout figure panels.

To better understand the basis for the improvement in sound location coding with arousal in AC, we related the sound coding of individual neurons (z-scored decoding performance) to the contribution of pupil size to their activity (from the encoding model in Figure 5). In AC, sound coding had a weakly negative relationship with pupil modulation (Pearson’s correlation = -0.03), where the neurons with the strongest sound coding tended to have the weakest pupil modulation. In PPC, on the other hand, sound coding and pupil modulation were positively correlated (Pearson’s correlation = 0.17), as the same neurons can be modulated by both sound and arousal state in PPC. We then examined the modulation of neural population activity by sounds and by arousal state, in population activity space, where each dimension (axis) is the activity of one neuron and a population of n neurons has n dimensions (axes) (Cunningham and Yu, 2014). We found the direction of the ‘sound location axis’, along which left vs right sound locations were best distinguished in this n-dimensional space, and the direction of the ‘pupil axis’, along which pupil size best explained population activity, in each AC and PPC population (See Materials and Methods, 11.1). We then measured the angle between these axes. In AC, these angles were more orthogonal to each other than in PPC (p=0.0006).

The correlation structure of population activity influences the amount of information that can be encoded across neurons (Averbeck et al., 2006; Panzeri et al., 2022). Specifically, the slope of the relationship between signal and noise correlations determines whether noise correlations limit the information that can be encoded by the population. In other words, greater shared variability among neurons with similar tuning preferences reduces the information that the population can encode. As a consequence, reductions in noise correlations among neuron pairs with similar tuning preferences or increases in noise correlations among neuron pairs with opposing tuning preferences would both theoretically increase the encoding capacity of a neural population (Averbeck et al., 2006; Panzeri et al., 2022). We then wondered whether transitions between arousal states also induced different changes in the correlation structure in AC and PPC, contributing to the improvements in sound location coding in AC. We compared pairwise noise correlations in high and low arousal conditions for both regions, and sorted these by the signal correlations between neuron pairs. Pairwise noise correlations among neurons with similar sound location preferences (high signal correlations) were reduced in the high arousal state in AC (p=0.002). In PPC, on the other hand, these similarly tuned neurons instead became more correlated with arousal (p<0.0001). In neuron pairs with negative signal correlations (different tuning), noise correlations were enhanced in AC (p = 0.0015), but did not change with arousal in PPC (p = 0.235). Taken together, the arousal-induced changes in the correlation structure of population activity suggest that the improvement in sound location coding in AC could result from a reduction in shared variability across neurons with similar sound preferences, in addition to the generalized suppression in sound-evoked responses during locomotion (Schneider et al., 2014; Bigelow et al., 2019; Yavorska and Wehr, 2021) that could sharpen response selectivity.

## Discussion

Our goal in this study was to determine whether the activity of inhibitory interneurons is differently modulated by behavioral state across the cortical processing hierarchy. We have measured spontaneous and sound-driven activity in populations of somatostatin-expressing inhibitory interneurons (SOM) and Non-SOM neurons in layer 2/3 of auditory cortex (AC) and in posterior parietal cortex (PPC), while mice transitioned between arousal states.

In the absence of sound stimulation, the effects of arousal state on spontaneous activity in AC were complex. Although heightened arousal had a slight positive effect on activity in AC at the population level, activity in individual AC neurons could be positively or negatively modulated, as has been described previously by others (Bigelow et al., 2019; Yavorska and Wehr, 2021). Short spurts of running, as we observed in our imaging sessions (Figure S1D), are associated with a net depolarization in AC (Shimaoka et al, 2018), which likely contributes to the net positive relationship between spontaneous AC activity and running observed here. In contrast, the spontaneous activity of most PPC neurons increased during these heightened arousal states, and SOM neurons were even more strongly and uniformly modulated than Non-SOM neurons.

The behavioral state transitions in our study involved increases both in pupil size and in locomotion (Figure 4A). It is thus crucial to emphasize that the effects of heightened arousal state on neural activity described here include a mixture of motor-related and ‘true’ arousal-related effects, such as those resulting from increases in norepinephrine and acetylcholine release within AC and PPC. While locomotion, whisking, and pupil dilations have all been considered as behavioral correlates of an animal’s arousal state, motor-related feedback acts on specialized circuits within visual, somatosensory, and auditory cortices, determining whether motor behavior positively or negatively influences neural activity (Lee et al., 2013; Fu et al., 2014; Schneider et al., 2014; Pakan et al., 2016; Bigelow et al., 2019; Yavorska and Wehr, 2021). This motor-related feedback acts in concert with neuromodulatory inputs more directly related to the animal’s arousal state, leading to state-dependent network changes that are specialized across cortical regions. It is unlikely that any aroused state is without a related change in fidgeting, facial movements, or other motor outputs (Musall et al., 2019; Stringer et al., 2019), and so the relative contributions of motor-related vs neuromodulatory inputs to neural activity must be defined for each brain region for a full consideration of state-dependent processing across the cortex. Here, we were able to disentangle the *unique* contributions of running velocity and pupil size to the activity of individual neurons in AC and PPC using an encoding model (Figure 5). In both regions, pupil size and running velocity had distinguishable contributions to the activity of SOM and Non-SOM neurons. The magnitude of unique pupil contributions was similar across all groups, but running contributions to activity were significantly stronger in PPC, particularly among SOM neurons.

In AC, the effects of arousal and locomotion can oppose each other, as motor-related feedback activates PV neurons, reducing sound-evoked responses during locomotion (Schneider et al., 2014; Bigelow et al., 2019; Yavorska and Wehr, 2021). We observed a small number of Non-SOM neurons in AC that were strongly and positively modulated with locomotion, which may include the PV neurons mediating the locomotion-related reduction of activity in other neurons (Schneider et al., 2014). In PPC, motor-related effects were largely positive, and were particularly strong among SOM neurons, suggesting that, unlike in AC (Schneider et al., 2014), motor feedback does not target PV neurons in PPC. Instead, SOM neurons in PPC may inhibit PV neurons during locomotion, disinhibiting excitatory neurons and further enhancing the arousal-related increase in activity in the local population. It will be interesting to determine whether this is the case in future studies. Recently, we also discovered that activity within the SOM population is highly coordinated, especially in PPC (Khoury et al., 2022). As a result, the transitions from low to high arousal states would trigger highly coordinated SOM population events, which could strongly impact the local network activity state (Chen et al., 2015; Veit et al., 2017; Wang and Yang, 2018). In the future, causal manipulations of SOM neurons, mimicking their activity during arousal transitions, will help reveal the impact of these coordinated SOM activity events on the activity and coding in the local population.

The effects of arousal on stimulus coding have been examined in depth across primary sensory cortices (Zhou et al., 2014; McGinley et al., 2015; Vinck et al., 2015; Shimaoka et al., 2018; Lin et al., 2019). Previous studies revealed that moderate levels of arousal optimally impact sensory coding (McGinley et al., 2015; Lin et al., 2019), aligning well with the Yerkes-Dodson (inverted-U) relationship between arousal and perceptual task performance (Waschke et al., 2019). Unlike these studies, we did not observe a decrement in sound location coding in the highest arousal state, instead measuring a modest improvement in decoding accuracy in the high arousal state. Mice in our experiments were most likely to visit two separable behavioral states (stationary/unaroused or running/aroused), without the gradation of different levels of arousal observed by others. As a result, the “high” arousal state described here likely includes a mixture of the moderate and high arousal states defined by others (McGinley et al., 2015). Because sensory coding in PPC depends on the behavioral relevance of stimuli (Fitzgerald et al., 2011; Pho et al., 2018), we expected that sound coding would also improve in PPC with heightened arousal, when mice might be more aware of the sound stimuli. Surprisingly, despite the more pronounced positive modulation of activity in PPC that accompanied increases in arousal and locomotion (Figure 4), the behavioral state did not affect sensory encoding in PPC (Figure 6). Our results imply that arousal alone is not sufficient to improve sensory coding in PPC outside of a task context, supporting the idea that task engagement and arousal modulate sensory responses through separate pathways (Saderi et al., 2021).

Finally, to better understand the basis of the improvement of sound coding with arousal in AC (and lack thereof in PPC), we related sound coding and pupil-related effects on population activity and its correlation structure. In AC, sound coding and pupil modulation were strongest in distinct sets of neurons (Figure 6F), and with heightened arousal, shared variability was reduced among neurons with similar tuning (Figure 6J). The net result was an improvement in our ability to decode sound information from AC population activity in the heightened arousal state, as the responses of the population to different stimuli were more separable (schematized in Figure 6L). In PPC, on the other hand, sound coding and pupil modulation were inter-mixed within the same neurons (Figure 6G), and shared variability *increased* among neurons with similar tuning (Figure 6K). As a consequence, although activity was strongly modulated in PPC with arousal, the separability of population responses to different sounds was not affected (Figure 6M). As has been recently reviewed, positive noise correlations among neurons with similar tuning limit the information that a population can encode (Averbeck et al., 2006; Panzeri et al., 2022), which is consistent with our results. To summarize, arousal had different effects on the correlation structure of population activity in AC and PPC. In AC, the result was better separability of the population responses to different sounds, while in PPC the effects of arousal on population activity were information-limiting (Averbeck et al., 2006; Panzeri et al., 2022).

To conclude, we have characterized the effects of the global arousal state on population activity in sensory and association cortex, by measuring neuronal activity during fluctuations in arousal and locomotion. In AC, but not PPC, sensory representations were enhanced with arousal, even when not behaviorally relevant. An important future direction will be to determine whether global shifts in arousal affect the coding of behaviorally relevant information in PPC, and whether local inhibitory circuits can provide a gating mechanism to enhance PPC’s encoding of behaviorally relevant sensory information.

## Acknowledgments

We thank Chengcheng Huang, Ross Williamson, and members of the Runyan lab for comments on the manuscript. This research was supported in part by the University of Pittsburgh Center for Research Computing through the resources provided. Judith Joyce Balcita-Pedicino performed histology and immunohistochemistry. We thank the GENIE project (Janelia) for making GCaMP sensors available for use. We thank the developers of Suite2P and Wavesurfer. This work was supported by the Andrew W. Mellon Predoctoral Fellowship, NIH Predoctoral Training Grant in Basic Neuroscience (T32 NS007433-21), Pew Biomedical Scholars Program, the Searle Scholars Program, the Klingenstein-Simons Fellowship Award in Neuroscience, and NIH grants NIMH DP2MH122404, NINDS R01NS121913.

## Supplementary Figures

**Figure S1:**
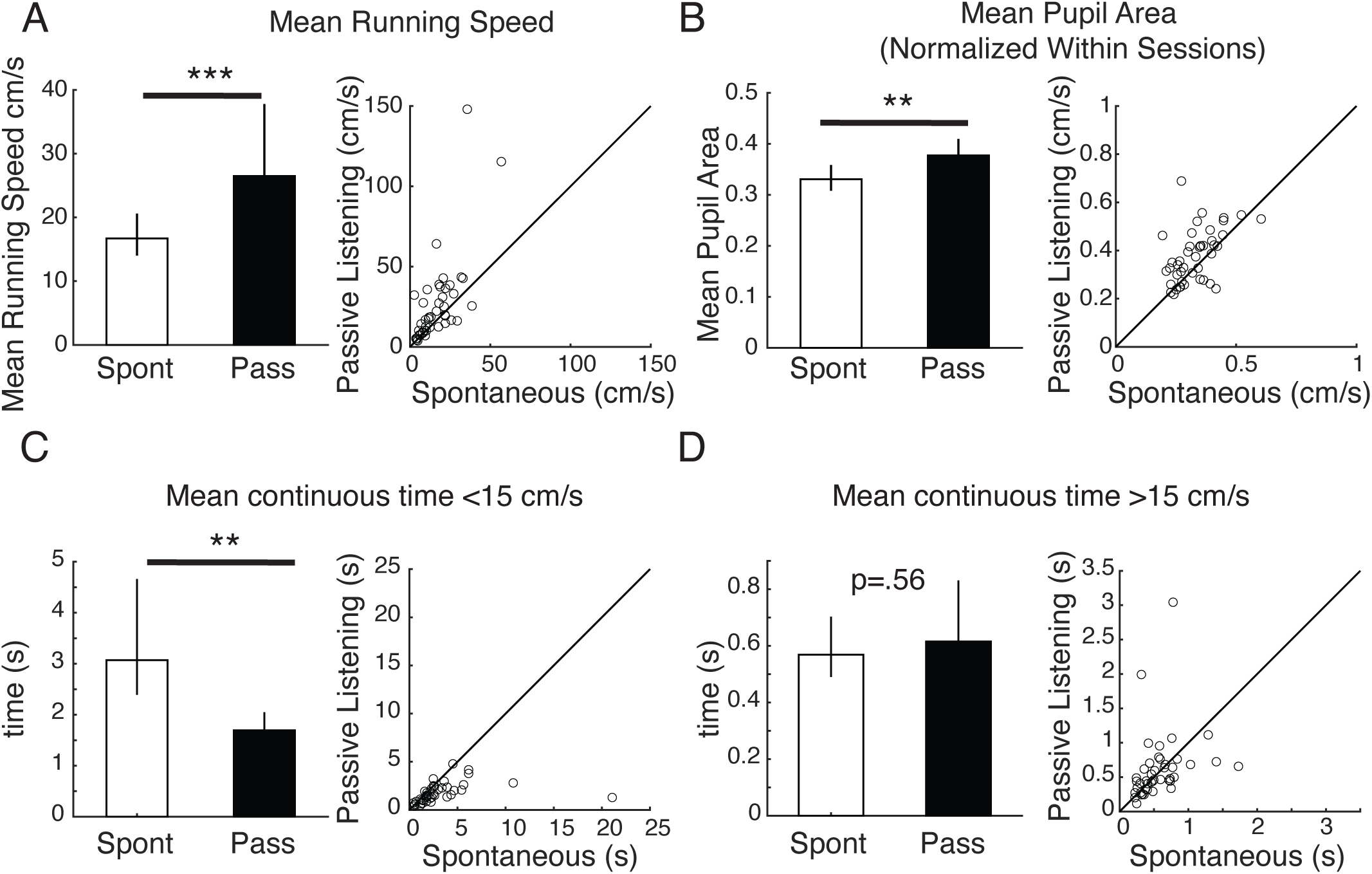
Characterizing running behavior and pupil size across contexts. (A) Left: Mean running speed in the spontaneous and passive listening behavioral contexts. Right: Mean running speed during the passive listening context plotted against the mean running speed during the spontaneous context, for each imaging session, p=.0003. (B) Left: Mean, maximum normalized pupil area during the spontaneous and passive listening contexts. Right: Mean pupil area in the two contexts for each imaging session. p=.002. (C) Left: Mean time spent running less than 15 cm/s in spontaneous and passive listening contexts. Right: Time spent running less than 15 cm/s during the two contexts, for each imaging session, p<.0001. (D) Left: Mean time spent running faster than15 cm/s in spontaneous and passive listening contexts. Right: Time spent running faster than 15 cm/s in the two contexts, for each imaging session. N=44 imaging sessions for all panels. Error bars indicate bootstrapped 95% confidence intervals, ***p<0.001; **p<.01.

**Figure S2:**
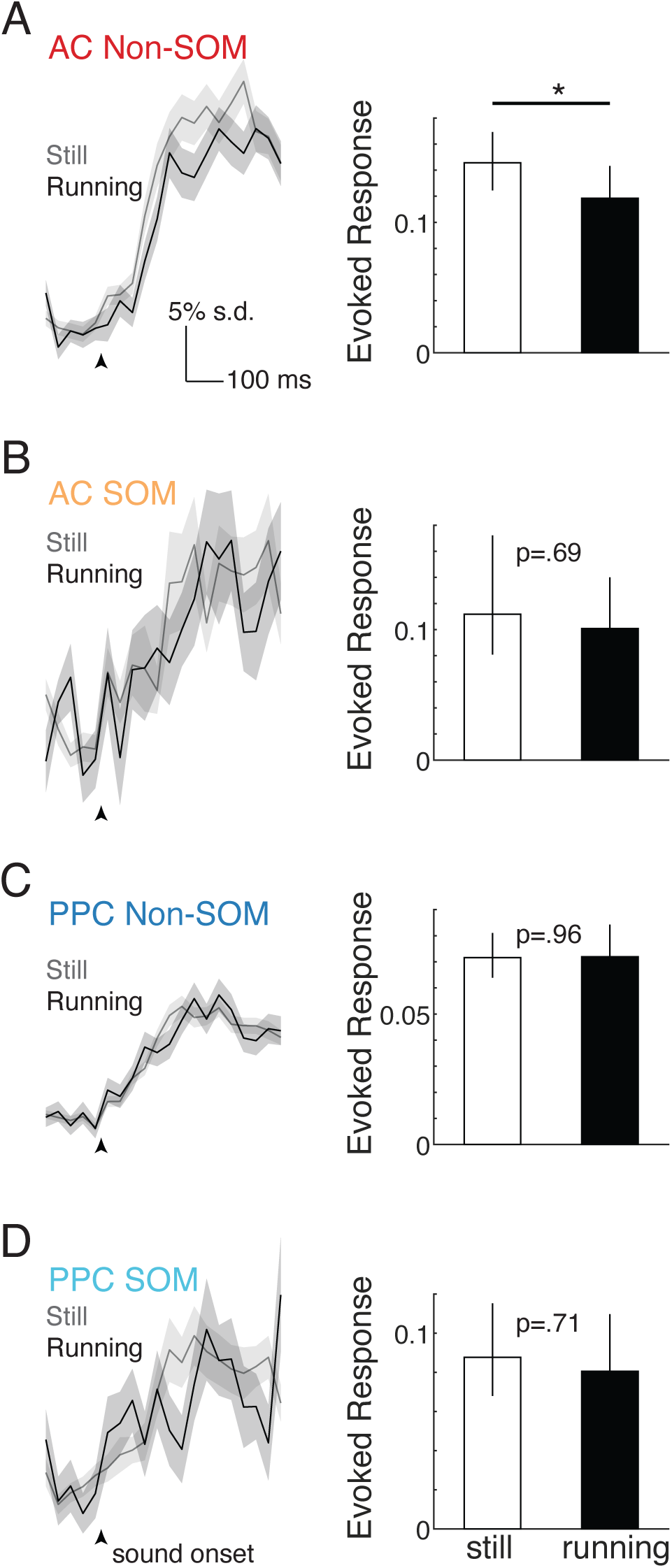
Sound evoked responses during periods of stillness and running. (A) Left: Sound-evoked activity of sound-responsive AC Non-SOM neurons (See: Figure 1K, Materials and Methods), in response to the preferred sound location. Gray: Responses during trials when mice were running less than 15 cm/s. Black: Responses during trials when mice were running faster than 20 cm/s. Sound-evoked activity was baseline-subtracted using the preceding 240 ms window for ease of display. Black arrow indicates sound onset. Right: Mean sound-evoked activity during the first 500ms of sound presentation during still and running periods, n=432, p=0.017 (paired permutation test with 10,000 iterations here, and throughout). Subsampling to match locomotion condition and repetition was repeated 10x for each sound location. (B) As in A, for AC SOM neurons, n=59 (C) As in A-B for PPC Non-SOM neurons, n=901 (D) As in A-C for PPC SOM neurons, n=112, *p<0.05.

